# Metabolite Damage and Damage-Control in a Minimal Genome

**DOI:** 10.1101/2021.12.01.470718

**Authors:** Drago Haas, Antje M. Thamm, Jiayi Sun, Lili Huang, Lijie Sun, Guillaume A.W. Beaudoin, Kim S. Wise, Claudia Lerma-Ortiz, Steven D. Bruner, Marian Breuer, Zaida Luthey-Schulten, Jiusheng Lin, Mark A. Wilson, Greg Brown, Alexander F. Yakunin, Inna Kurilyak, Jacob Folz, Oliver Fiehn, John I. Glass, Andrew D. Hanson, Christopher S. Henry, Valérie de Crécy-Lagard

**Affiliations:** Department of Microbiology and Cell Science, University of Florida, Gainesville, FL 32611, USA; Horticultural Sciences Department, University of Florida, Gainesville, FL 3261, USA; Food Science and Human Nutrition Department, University of Florida, Gainesville, FL 32611, USA; J. Craig Venter Institute, La Jolla, CA 92037, USA; Chemistry Department, University of Florida, Gainesville, FL 32611, USA; Maastricht Centre for Systems Biology (MaCSBio), Maastricht University, 6200 MD Maastricht, The Netherlands; Department of Chemistry, University of Illinois at Urbana-Champaign, Urbana, IL 61801, USA; Department of Biochemistry and the Redox Biology Center, University of Nebraska, Lincoln, NE 68588, USA; Department of Chemical Engineering and Applied Chemistry, University of Toronto, Toronto, ON M5S 3E5, Canada; Centre for Environmental Biotechnology, School of Natural Sciences, Bangor University, Bangor, LL57 2UW, UK; West Coast Metabolomics Center, UC Davis, Davis, CA 95616, USA; Data Science and Learning, Argonne National Laboratory, Argonne, IL 60439, USA; Consortium for Advanced Science and Engineering, The University of Chicago, Chicago, IL 60637, USA; University of Florida Genetics Institute, Gainesville, FL 32611, USA11

**Author notes:** Sanofi, 13 Quai Jules Guesde Vitry-sur-Seine 94400, France (Drago Haas). Havas Life Bird and Schulte, Urachstrasse 19, 79102 Freiburg im Breisgau, Germany (Antje M. Thamm). Captozyme, 1622 NW 55th Place, Gainesville, FL 32653, USA. Lingnan Medical Research Center, Guangzhou University of Chinese Medicine, Guangzhou, Guangdong, China, 510006 (Lili Huang). Ginkgo Bioworks, 27 Drydock Ave 8th Floor, Boston, MA 02210. Corresponding Authors: Valérie de Crécy-Lagard and Christopher S. Henry. Possible reviewers* Carole Linster Vadim Gladyshev. Zoran Nikoloski https://www.mpimp-golm.mpg.de/13193/Zoran_Nikoloski. Pedro Mendes http://www.comp-sys-bio.org/pedro/Mendes.html. Paco Baroma-Gomez https://langebio.cinvestav.mx/en/Dr-Francisco-Barona.

## Abstract

Analysis of the genes retained in the minimized Mycoplasma JCVI-Syn3A genome established that systems that repair or preempt metabolite damage are essential to life. Several genes with known metabolite damage repair or preemption functions were identified and experimentally validated, including 5-formyltetrahydrofolate cyclo-ligase, CoA disulfide reductase, and certain hydrolases. Furthermore, we discovered that an enigmatic YqeK hydrolase domain fused to NadD has a novel proofreading function in NAD synthesis and could double as a MutT-like sanitizing enzyme for the nucleotide pool. Finally, we combined metabolomics and cheminformatics approaches to extend the core metabolic map of JCVI-Syn3A to include promiscuous enzymatic reactions and spontaneous side reactions. This extension revealed that several key metabolite damage-control systems remain to be identified in JCVI-Syn3A, such as that for methylglyoxal.

## Introduction

A foundational goal of synthetic biology was to create a minimal living organism by a bottom-up approach (1). This goal was reached in 2016 with the creation of JCVI-Syn3.0 (2). This organism, based on the blueprint of the ruminant pathogen *Mycoplasma mycoides capri* serovar LC GM12, a Gram-positive bacterium, was built by combining DNA synthesis, recombination, and genome transplantation techniques, and was designed to include only genes that are required for survival or to support a reasonable growth rate (428 protein-coding genes and 34 genes for RNAs) (2). The initial strain JCVI-Syn3.0 was extremely fragile and another derivative with an additional 18 genes, JCVI-Syn3A was found to be more stable and was the basis for the recently published metabolic model (3). Surprisingly, at the time of publication in 2016, ∼30 % of the genes in JCVI-Syn3.0 could not be assigned a specific function. The initial annotation has since been improved by manual curation (4), through the creation of a detailed metabolic model (3) and through further in silico analyses (5) but ∼85 proteins with unknown or just broadly defined function remain (Supplemental data S1). These unknowns cannot all be missing parts of synthesis/breakdown pathways as the metabolic reconstruction only identified a few such gaps, namely four metabolic and eight transport reactions (3).

A crucial area of metabolism that is left out of classical metabolic models is metabolite damage and repair. Enzymes make mistakes and metabolites can undergo spontaneous chemical reactions (for classical examples see reference (6)). These types of uncontrolled metabolite damage are ever-present and, when the resulting side-products have toxic effects, their accumulation can impose a fitness cost (6, 7). In recent years, it has been shown that many enzymes of formerly unknown function repair or pre-empt metabolite damage (8), that human diseases are caused by mutations in metabolite repair enzymes (9–11), and that pathway engineering can fail unless the necessary repair enzymes are installed (12). The emerging recognition of the nature and extent of metabolite damage and repair raised the question of the importance of metabolite repair for the survival and growth of a minimal genome like JCVI-Syn3/3A. By combining expert manual curation, comparative genomics, metabolomics, metabolic modeling, chemoinformatics, and experimental validation, we identified a set of chemical damage reactions likely to occur in JCVI-Syn3 and some of the damage repair and preemption activities that are encoded (or predicted) by this minimal genome.

## Results and Discussion

### Identification and experimental validation of homologs of known metabolite repair enzymes

To identify the metabolite repair enzymes in JCVI-Syn3A we first manually scanned its proteome for homologs of known metabolite repair enzymes (6, 12, 13)(see Supplemental data S1 and supplemental methods). Several were found, as follows.

#### 1. 5-FCL

5-Formyltetrahydrofolate (5-CHO-THF) is a by-product of serine hydroxymethyltransferase (SHMT) (14)(Fig. 1A) that inhibits folate-dependent enzymes and must therefore be removed from the folate pool (15). Of various enzymes known to recycle 5-CHO-THF (16), the most widespread is 5-formyltetrahydrofolate cyclo-ligase (5-FCL) (encoded by the *fau/ygfA* gene (16) in *E. coli*). The JCVI-syn3A genome encodes a 5-FCL homolog (JCVISYN3A_0443; 24% identity over 93% coverage); this gene was confirmed to encode an active 5-FCL by a complementation assay. Specifically, an *E. coli* K12 Δ*ygfA* strain does not grow on M9 minimal medium with 0.2% glucose as carbon source and 20 mM glycine as sole nitrogen source (16)(Fig. 1B). Expression of JCVISYN3A_0443 from a pUC19 derivative plasmid allowed complementation of the growth phenotype (Fig. 1B). Note that the essentiality of JCVISYN3A_0443 might not be due to the repair function alone, but also to a role in 5-CHO-THF-polyglutamate salvage as a unique source of 5,10-methenyltetrahydrofolate-polyglutamate (3).

**Figure 1.**
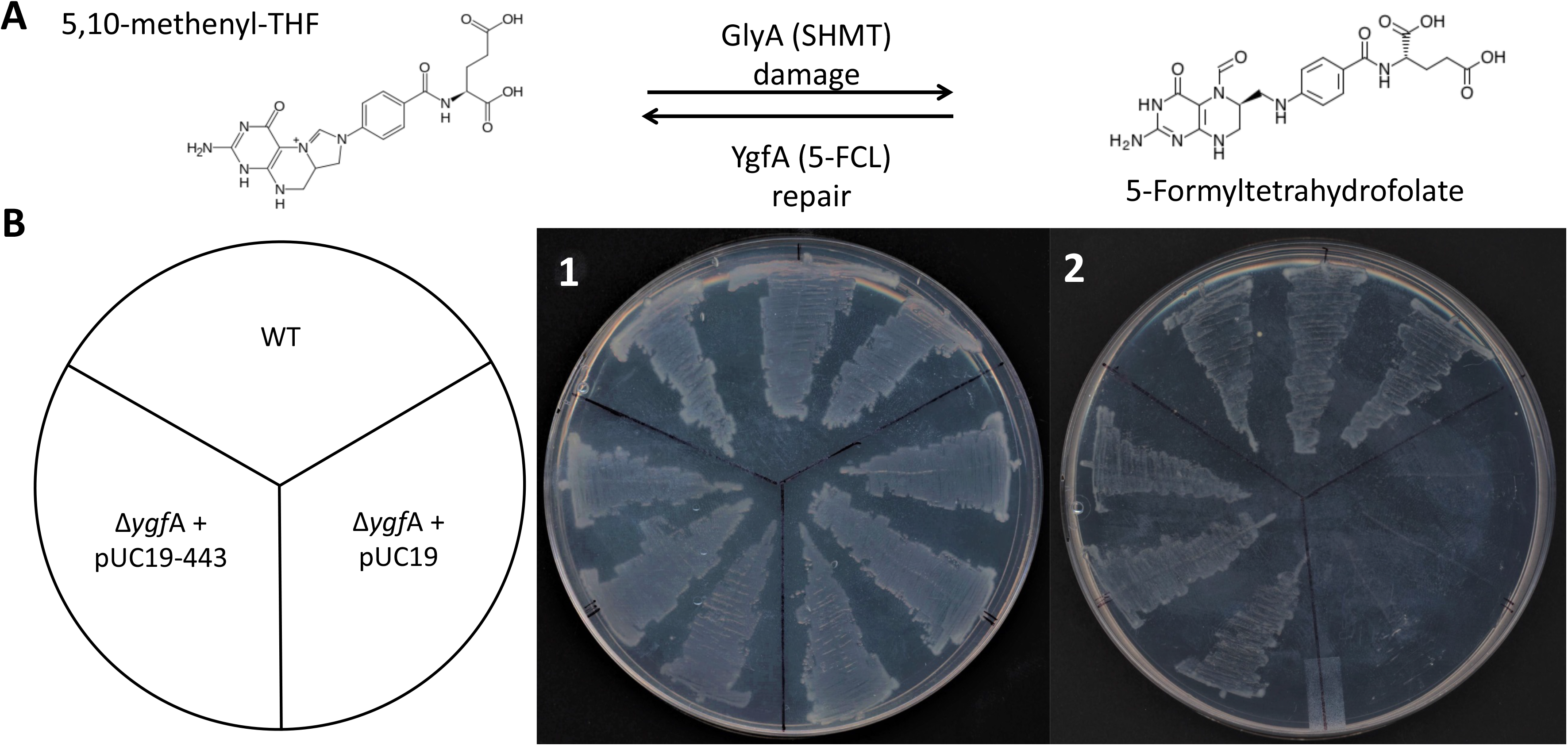
5-FCL activity is encoded by JCVI_0400. **(A)** Enzymatic source and repair of 5-CHO-THF. **(B)** Growth phenotype of a WT *E. coli* BW25113, Δ*ygf*A mutant and, Δ*ygf*A mutant expressing JCVI_0443 gene on M9 minimal medium (0.4% glucose) with (**1)** 20 mM NH_4_Cl or (**2)** 50 mM glycine as sole nitrogen source. Plates were incubated for 3 days at 37°C.

#### 2. Cellular thiol reductases

Like all organisms grown in the presence of oxygen, JCVI-syn3A will encounter oxidative stress that can damage macromolecules. Maintaining protein and small-molecule thiol groups in their reduced state is critical for cellular redox homeostasis (17). Thioredoxin/thioredoxin reductase is the dominant protein thiol oxidoreductase system in many organisms, using reducing equivalents ultimately derived from NAPDH (18, 19). The JCVI-Syn3A genome encodes homologs of the thioredoxin system proteins (TrxB/JCVISYN3A_0819 and TrxA/JCVISYN3A_0065) that are most likely involved in reducing disulfide bonds in proteins and have already been partially characterized in other *Mycoplasma* species (Fig. 2A)(20, 21). Both genes are essential (Supplemental data S1), supporting a key role of the TrxA/TrxB system in disulfide bond reduction. Note, however, that thioredoxin is also the hydrogen donor to ribonucleotide reductase, and thus JCVISYN3A_0819 and JCVISYN3A_0065 may be essential due to this possible connection to deoxyribonucleotide biosynthesis (20, 22).

**Figure 2.**
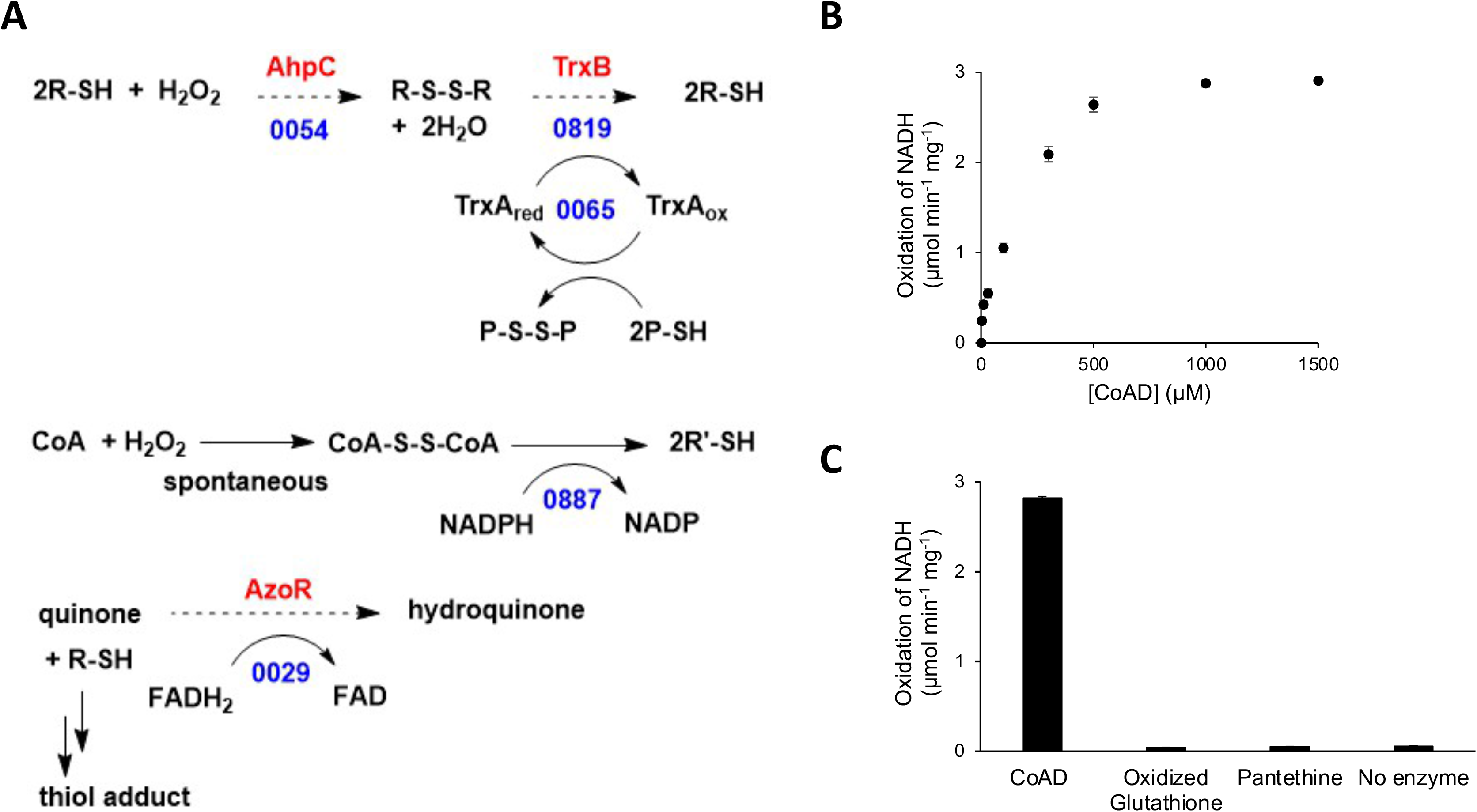
Predicted and validated redox buffering systems in JCVI-Syn3. **(A)** Candidates for H_2_O_2_ detoxification systems of JCVI3, experimentally validated are in solid arrows, only the number of the locus tags are given, P is for protein, R is for small molecule. (**B)** CoADR Michaelis-Menten saturation curve for the determination of the Km and kcat for CoAD consumption. **(C)** CoADR is specific towards oxidized CoA with no activity towards other tested disulfides

JCVI3_0887 was found to be a homolog of CoA disulfide reductase (CoADR), which is proposed to be the major mechanism to maintain redox balance in certain bacteria (23). Because CoA is required for several reactions in the JCVI-syn3A metabolic model and is predicted to be imported from the medium, CoADR could be a minimalist solution to detoxify H_2_O_2_. We therefore tested the CoADR activity of the JCVISYN3A_0887 protein *in vitro*. JCVISYN3A_0887 was found to be an active CoAD reductase that operates well at physiologically relevant pH (pH 7.5) (24) with reasonable *K*_M_ (0.17 mM) and *k*_cat_ (2.8 s^-1^) values (Fig. 2B). It had no detectable activity against oxidized glutathione or pantethine (Fig. 2C). While we cannot eliminate the possibility that reduced glutathione is imported from the medium and oxidized glutathione is exported, this is a less parsimonious solution to the redox balance problem than the CoA-based solution above.

### Functional analysis of orphan HAD family proteins identifies a nucleotide phosphatase with possible dual roles

Our second strategy to identify metabolite repair enzymes was based on the fact that various hydrolases of uncertain or unknown function were subsequently shown to participate in metabolite repair (8)Five genes encoding stand-alone members of the HAD (<u>h</u>alo<u>a</u>cid <u>d</u>ehalogenase) hydrolase family (25) were identified in the JCVI-Syn3A genome (Supplemental data S1) and are conserved in the recently analyzed close mollicute relative *Mesoplasma florum* L1 (26) (Table 1). Such HAD hydrolases often participate in metabolite repair or homeostasis, as many damaged and/or toxic intermediates are phosphorylated (e.g. phosphosugars), and the first step in their recycling or removal requires a phosphatase (8, 27).

**Table 1.** Members of the HAD family of unknown function encoded by JCVI-Syn3

| Gene | Family | Essential | Best 3 substrates<br>Activity *in vitro* * | Physical clustering | *M. florum* ortholog<br>locus tag and<br>essentiality** |
| --- | --- | --- | --- | --- | --- |
| JCVISYN3A_0066 | Cof subfamily of IIB<br>subfamily of HAD<br>superfamily | no | pNPP, FMN,<br>CoA | Between 5S rRNA gene<br>and thioredoxin | Mfl169 (NE) |
| JCVISYN3A_0077 | Cof-like hydrolase, HAD<br>superfamily | no | Fru-1P, Ery-4P | Between *tsaD* and *aspS* | Mfl614 (E) |
| JCVISYN3A_0710 | Cof subfamily of IIB<br>subfamily of HAD<br>superfamily | yes | Could not clone | Between tRNA genes<br>and predicted<br>phosphonate transporter<br>genes | Mfl513 (E) |
| JCVISYN3A_0728 | HAD superfamily<br>hydrolase subfamily IIB,<br>protein | no | GMP<br>XMP<br>2-deoxy-glucose-6P | Between glycolysis<br>genes | Mfl503 (E) |
| JCVISYN3A_0907 | Cof-like hydrolase, HAD<br>superfamily | no | N-acetyl-D-glucosamine-6P<br>Fructose-1P<br>N-acetyl-D-glucosamine-1P | Between YidC and<br>choline kinase-like | Mfl680 (NE) |

\*Abbreviations in Table S1; \*\* (E), essential; (NE)=non-essential in *M. florum*

Comparative genomic analysis of the stand-alone HADs did not point to clear functional hypotheses, except for JCVISYN3A_0728, whose location in a predicted operon with triose-phosphate isomerase and phosphoglycerate mutase suggested a role in sugar phosphate metabolism (Table 1). Possible functions for the HAD family proteins included: 1) metabolite repair enzymes on substrates to be identified; 2) missing phosphatases involved in primary metabolism identified by the metabolic model such as sedoheptulose 1,7-bisphosphate phosphatase or phosphatidate phosphatase; 3) nucleotide phosphatases involved in dNTP pool maintenance. To discriminate among these hypotheses, we combined biochemical, genetic, and metabolomics studies.

All four of the HAD proteins that we were able to express in *E. coli* (JCVISYN3A_0066, JCVISYN3A_0077, JCVISYN3A_0728, JCVISYN3A_0907) were tested for activity against a panel of 94 phosphatase substrates (Supplemental Table S1) (28). The four proteins had detectable activity against the general phosphatase substrate *p*-nitrophenyl phosphate (*p*NPP) and different sets of physiological substrates (Fig. S1). The JCVISYN3A_0728 enzyme was active against a wide range of nucleoside and sugar phosphates, the JCVISYN3A_0907 and JCVISYN3A_0077 enzymes were active against narrower ranges of sugar phosphates, and the JCVISYN3A_0066 enzyme was active against FMN and CoA. That sugar phosphates are among the best substrates of the JCVISYN3A_0728 enzyme is consistent with the genomically-predicted role in sugar phosphate metabolism, but no specific function or substrate could be assigned. Note, however, that the 94-substrate panel did not include damaged sugar phosphates.

We attempted to delete HAD-encoding genes in the JCVI-syn3A host as described in the Methods section. We expected this to be possible because transposon bombardment of the JCVI-syn3A genome indicated all five HADs were quasi-essential (i.e., required for fast growth but not essential for viability) [(3) and Supplemental data S1]. Deletion mutants were readily obtained for genes JCVISYN3A_0066, JCVISYN3A_0077, JCVISYN3A_0728, and JCVISYN3A_0907 (Supplemental data S2). Attempts to delete JCVISYN3A_0710 using two different methods were unsuccessful (Supplemental data S2). It could be that the deletion of this gene resulted in an extremely slow-growing strain that could not be recovered using these approaches, or that JCVISVN3A_0710 is in fact essential and the transposon insertions in the gene were artifacts. The fact that the same gene is also essential in *M. florum* (Table 1) would suggest that the latter hypothesis is correct

We observed no major differences in growth rates between JCVI_Syn3A and any of the HAD mutants (Fig. S2). To conduct a metabolomics analysis, the four mutants and the parental JCVI-Syn3A control were grown in SP4-KO medium and harvested at the same point of log-phase growth. Three biological replicates were prepared and three technical replicates from each biological sample were distributed and further pelleted/rinsed/flash frozen/stored. Untargeted metabolomic analysis of the mutant samples was carried out to screen for a broad range of possible metabolic disruptions. The extraction, detection, and analyses of the metabolites are described in detail in the Supplemental methods and Supplemental Tables S2 and S3. A total of 4152 features were detected in JCVI-Syn3A samples using hydrophilic interaction liquid chromatography (HILIC) and mass spectrometry (Supplemental data S3). Metabolites were annotated using accurate mass in addition to matching experimental MS/MS spectra to MS/MS library spectra (MS/MS match) and/or matching experimental peaks to an in-house accurate mass/retention time library (m/z-RT match). In total 522 metabolites were annotated as known metabolites in cultures of JCVI-syn3A and mutants of JCVI-syn3A. Of these annotated metabolites, 70 had both MS/MS and m/z-RT matches, 324 were annotated based on MS/MS matches, 100 were annotated based on m/z-RT matches, and 28 annotations had MS/MS matches linked to a small number of candidate compounds (e.g., “hexose-phosphate”, which could be multiple phosphorylated six-carbon sugars) (Supplemental data S3). Technical variance was assessed by measuring 43 internal standard compounds, which are non-endogenous chemicals added to each sample and had an average coefficient of variance of 8.9% (Supplemental data S3).

Partial least squares discriminant analysis was used to find the variable importance in projection (VIP) scores of each annotated metabolite. The fifteen metabolites with the highest VIP scores (Fig. 3 and Fig. S3) showed little contamination from media, as determined by chemical analysis of unused media along with mutant samples. Most of these metabolites were below the limit of detection in unused media, and most of the rest were present at much lower abundance in media than in samples (>30-fold higher in samples compared to media) suggesting little to no contamination from residual media in samples (Supplemental data S3). Two metabolites (cytidine and thiamine) were found at similar abundance in media as compared to samples, suggesting these two compounds may be influenced by media contamination.

**Figure 3.**
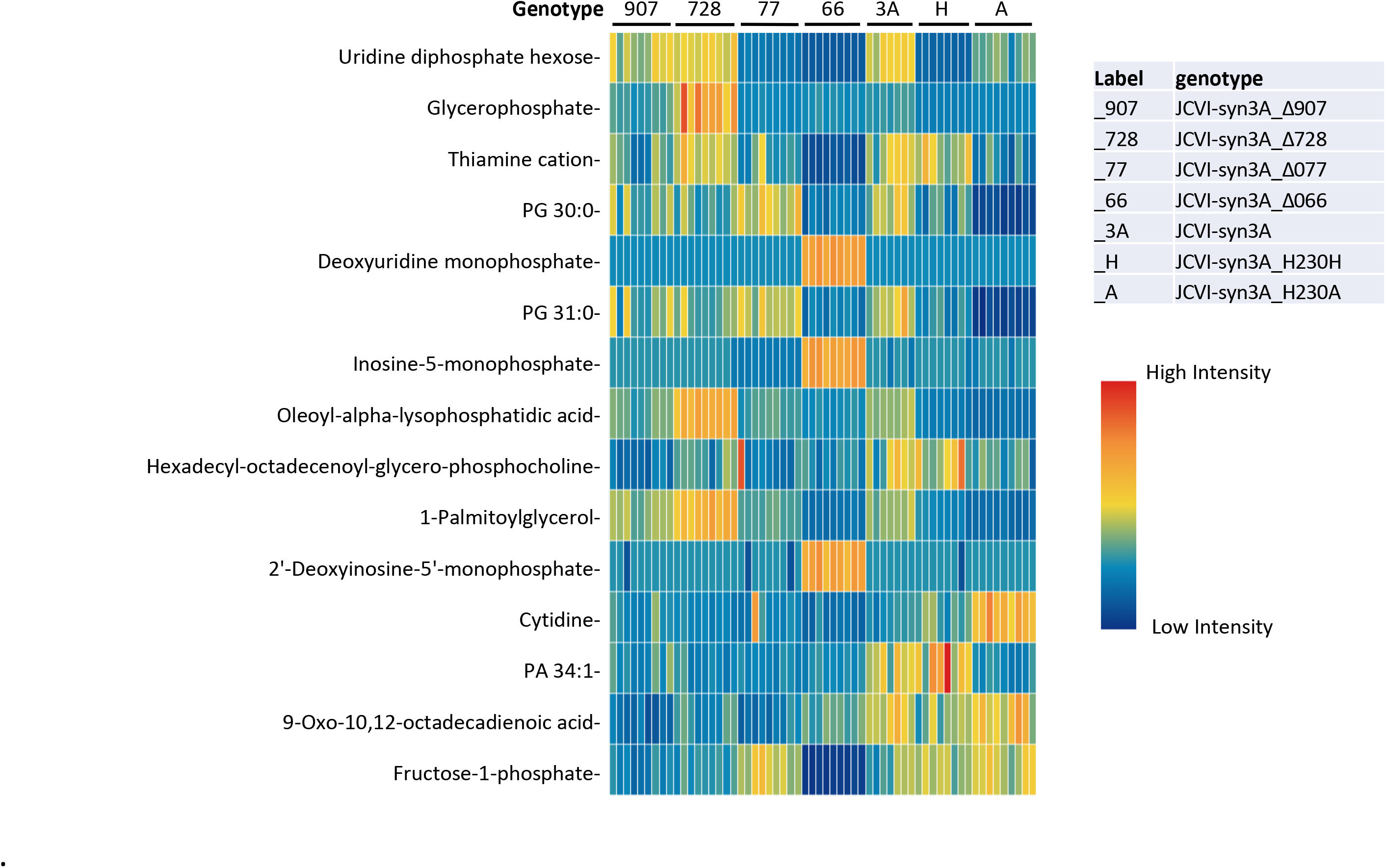
Heatmap including 15 metabolites from JCVI-syn3A mutant metabolomic analysis with highest VIP scores. Samples and genotypes are represented in columns. High intensity measurements as compared to average intensity are red/yellow, and low intensity measurements are represented by green/blue

Within this group of 15 metabolites with high VIP scores, the JCVISYN3A_0728 knockout showed significantly higher abundance of glycerophosphate, oleoyl lysophosphatidic acid, and palmitoylglycerol as compared to other genotypes (Fig. 3 and Fig. S3). We were not able to determine which form of glycerophosphate was increased, although the 3-phosphate is a priori more likely, being found in the metabolic model as a cardiolipin metabolism intermediate that is synthesized via phosphorylation of imported glycerol by GlpK (JCVISYN3A_0218). As alpha-lysophosphatidic acid is produced from glycerol phosphate (3), these results suggest that the JCVISYN3A_0728 hydrolase could be involved in hydrolyzing glycerol phosphate (or a derivative thereof).

The JCVISYN3A_0066 knockout had significantly higher deoxyuridine monophosphate, inosine monophosphate, and deoxyinosine monophosphate, and lower thiamine and fructose-1-phosphate abundance as compared to other mutants (Fig. 3 and Fig. S3). It is therefore possible that JCVISYN3A_0066 is one of the phosphatases previously found to hydrolyze the mononucleotides GMP, dAMP, dGMP, dUMP, and dTMP that were not identified in the metabolic reconstruction (3)and further biochemical characterization is needed.

### Comparative genomic approaches uncover a possible metabolite repair phosphatase

Comparative genomic analyses allowed us to propose a function for the YqeK HD family phosphohydrolase that is fused to nicotinic acid mononucleotide adenylyltransferase (NadD) in most mycoplasmas and strongly physically clustered with NadD in many other gram-positive organisms (29) (Fig. 4 and S4A). This strong association led us to propose that the YqeK protein repairs mistakes made by the NadD enzyme. The canonical activity of NadD is to adenylate nicotinate-ribonucleotide (NaMN) using ATP as a donor of the AMP moiety (Fig. 4A). However, if another NTP or the deoxy-form of ATP is used by mistake, this would create an erroneous intermediate that would need to be hydrolyzed. We therefore expressed the JCVISYN3A_0380 gene in *E. coli* as well as a variant encoding a His230Ala mutation (Fig. S4B). (The mutation of this residue, predicted to be critical for the phosphatase activity, was designed to stop product hydrolysis interfering with measurement of NadD activity.) The NadD protein of *Bacillus subtilis* was also expressed and purified as a benchmark. The mutant JCVISYN3A_0380 protein and *B. subtilis* NadD were then tested for *in vitro* activity with various nucleoside triphosphates as substrates. As shown Fig. 5A, we found that the adenylation activity of the JCVISYN3A_0380 His230Ala mutant was quite non-specific and actually worked better with dATP, CTP, or UTP than with the natural substrate, ATP, whereas the *B. subtilis* enzyme strongly preferred ATP. The JCVI-syn3 NadD enzyme can therefore readily form deoxy-adenosine, -cytidine, or -uridine analogs of the NAD precursor nicotinate adenine ainucleotide (NaAD) (which can presumably be converted to potentially inhibitory analogs of NAD and NADP).

**Figure 4.**
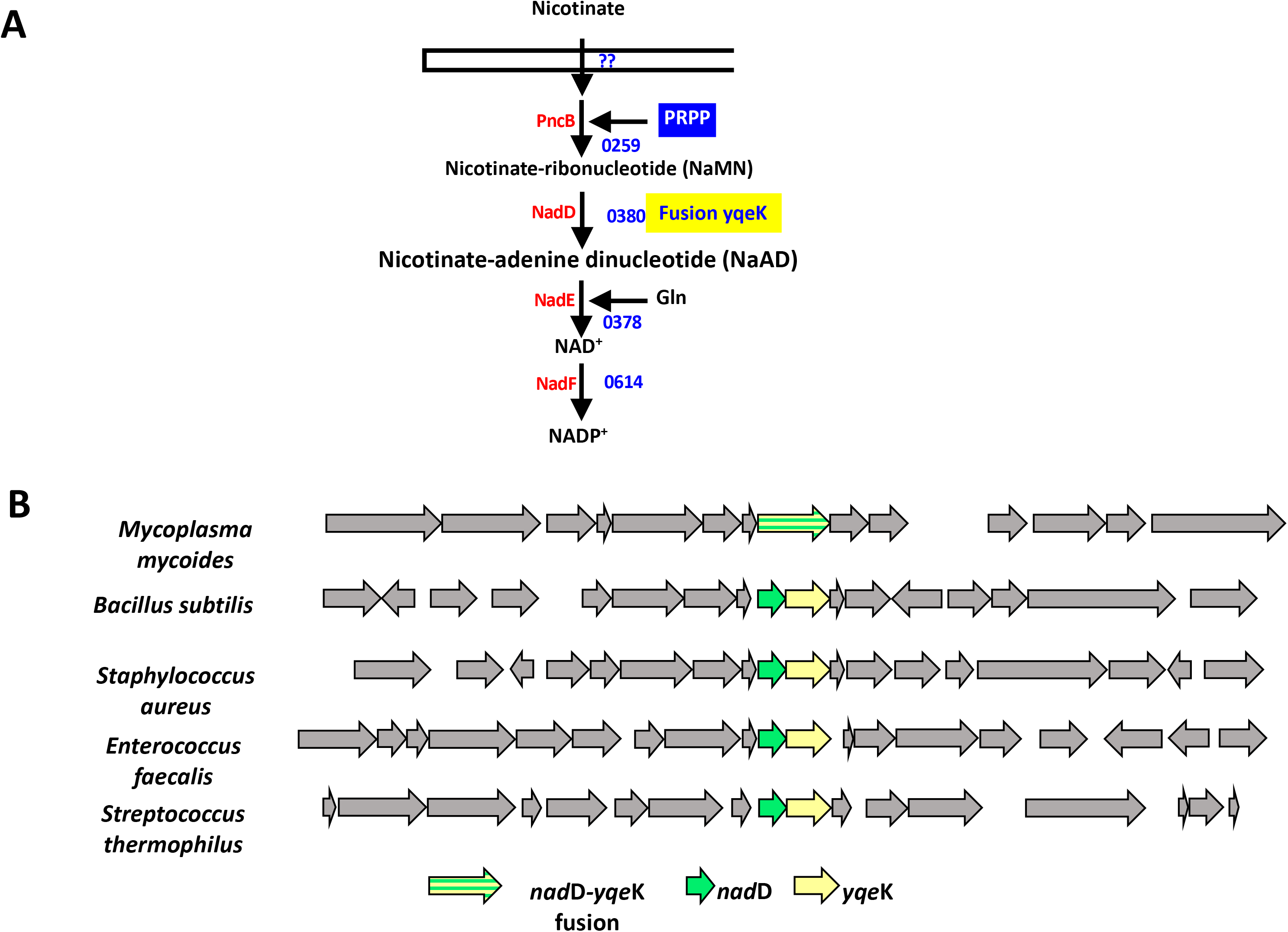
Predicted Hydrolase of unknown function is clustered or fused to NadD in many Firmicutes. **(A)** Predicted NADP^+^ synthesis pathway in JCVI-Syn3. **(B)** Physical clustering and fusions of *nadD* and *ykeK* homologs in several gram-positive Bacteria. The RefSeq identifiers for the *yqeK* genes used in descending order are: NP_975428.1, NP_390441.1, NP_372117.1, NP_816490.1, YP_140036.1. **(C)** Docked model of 2-deoxy-NaAD bound to the *C. acetobutylicum* YqeK (pdb code: 3CCG). The protein is shown in ribbon format (grey) with side chains as lines, two iron atoms are shown as spheres bound to the diphosphate of dNaAD. Tyrosine 82 (green) is modeled as two conformations in the crystal structure and forms a close interaction with the 2’carbon of dNaAD.

**Figure 5.**
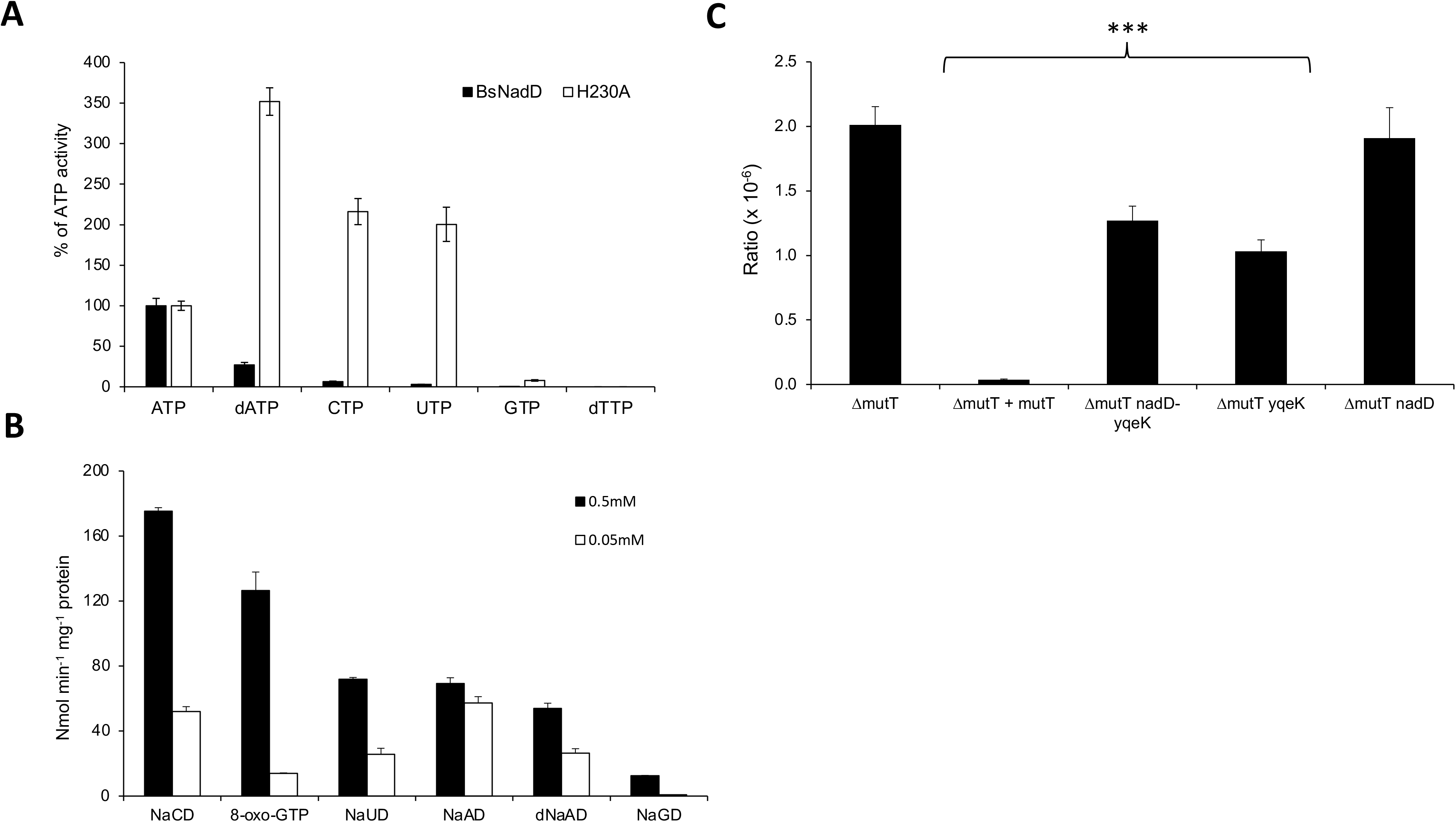
Biochemical analysis of the NadD and YqeK activities. **(A)** Relative reaction rates of *Bacillus subtilis* and JCVI syn3.0 NadD enzymes with NaMN and various nucleotides, calculated as percentage of the canonical reaction with ATP for each NadD enzyme. Enzymes were incubated with 2 mM NTP, 0.5 mM NaMN, 4 mM MgCl_2_ and 5 u/ml yeast inorganic pyrophosphatase for 5 min at 37° C. H230A has the conserved H in the active site of the YqeK domain mutated to ablate the HD activity and cleavage of nucleotides. **(B)** Activity of the expressed JCVI syn3.0 YqeK domain with different substrates. YqeK (0.2 μg) was incubated with 0.5 or 0.05 mM substrates, 1 mg/ml BSA and 2.0 mM MgCl_2_ for 20 min at at 37° C. Black bars are data for 0.5 mM substrates, white bars are data for 0.05 mM substrates. **(C)** Mutation ratio on LB rifampicin for strain Δ*mut*T with empty vector (pBAD24), Δ*mut*T with E. coli *mut*T in trans, Δ*mut*T with either the *nad*D-*yqe*K fusion gene JCVI_0380, or the *nad*D or *yqe*K domains alone. *** indicates a P-value <0.001 with experiments performed with four biological replicates and four technical replicates.

We then tested the JCVI-Syn3 YqeK domain for phosphatase activity using the different NaAD analogs that could be produced by the JCVI-syn3A NadD enzyme. As shown Fig. 5B, the YqeK domain has activity towards the cytosine (NaCD) and uracil (NaUD) analogs of NaAD that is as high or higher than that against NaAD itself (which agrees with the preference of the NadD domain to make these analogs).

In order to assess the relative binding capacity for predicted substrates for YqeK, we docked the proposed molecules into the structure. The active site was defined at the di-iron site and molecules were modeled with phosphate groups coordinated to the metals as seen in representative structures (PDB codes: CCG3, 2O08 and 2OGI). The conformations of the liganded substrates were optimized through rounds of energy minimization. The results show that the active site readily accommodates 2′-deoxy-NaAD (Fig. S4C) and interactions of the adenosine and nicotinate moieties are consistent with other NaAD-binding proteins (e.g., PDB 1NUQ and 2QTR). Of note, a conserved Tyr82 clashes with the 2′-hydroxyl of NaAD, not present in 2′-deoxy-NaAD.

In testing various possible substrates, we found the YqeK domain also had high activity against 8-oxo-GTP, although judging from relative activities with 0.05 mM and 0.5 mM substrate, the *K*_M_ is likely higher than for the other substrates tested (Fig. 5B). Consistent with this finding, we showed that the genes encoding the JCVI-syn3A NadD-YqeK fusion can partially complement the *E. coli mutT* high mutation rate phenotype (measured as Rif^R^ ratios) (Fig. 5C). The partial complementation was also observed when expressing the YqeK domain alone, but not the NadD domain alone. Finally, it was recently shown that YqeK of gram-positive bacteria such as *B. subtilis* or *M. pneumoniae* are members of a novel diadenosine tetraphosphate (Ap4A) hydrolase family (30). In combination, these observations suggest that YqeK is a versatile phosphatase with several functional roles. Unfortunately, the genetic dissection we performed in JCVI-Syn3A did not allow us to confirm any of these roles *in vivo*. Indeed, the available transposon insertion data ((3)and Supplemental data S1) suggested that the NadD domain is essential and the YqeK domain is quasi-essential as a few hits in the YqeK region of the gene were detected in the first Tn round, these disappeared after the fourth round of growth. We were unable to isolate a JCVISYN3A_0380 gene deletion mutant in JCVI-Syn3B despite several attempts. We were, however, able to construct a derivative that contains the His230Ala point mutation inactivating the YqeK activity (Supplemental data S2) that did not show any growth rate defect or any obvious metabolite imbalance (Fig. S2). Further studies will accordingly be needed to define the role of the YqeK proteins in metabolism.

### Metabolomics-driven exploration of damage and repair chemistry in JCVI-Syn3

Thus far, all of our damage and repair cases began with an analysis of specific classes of genes in the JCVI-Syn3A genome, and from these cases we see clear instances of metabolite damage and repair occurring in the JCVI-Syn3A strain. But do these examples represent isolated exceptions, or are they the tip of an iceberg of uncharacterized metabolic chemistry occurring even in the simplest organism that can currently be constructed? To gain insights into this question, we applied a more systematic exploratory approach that started with the metabolomics data generated from our JCVI-Syn3A cell samples (see Supplemental Table S3).

We focused this analysis specifically on the set of 480 metabolites observed in these samples that satisfied two criteria: (1) the mass-spectrum-observed metabolite was confidently identified with a fully defined molecular structure; and (2) the metabolite was at least as abundant in the JCVI-Syn3A cells as in the growth medium. Supplemental Table S4E contains this list of 480 metabolomics peaks filtered from the full set of metabolomics data provided in Table S3. We next compared the 480 identified peaks to the 33,978 compounds in the ModelSEED database (31), which includes all of KEGG (32) and MetaCyc (33), resulting in 217 (45%) matches (see Supplemental Table S4E). We next compared the 480 identified peaks to the 33,978 compounds in the ModelSEED database (32), which includes all of KEGG (33) and MetaCyc (34), resulting in 217 (45%) matches (see Supplemental Table S4E). This analysis revealed that over half the observed metabolites fall outside current biochemistry databases, and that even for compounds that do occur in existing databases, they take part in pathways that are not included in the current representation of JCVI-Syn3 metabolism. To predict potential chemical routes to as many of the observed metabolites as possible without limiting our search to known chemistry or straying too far from known JCVI-Syn3 metabolism, we applied the cheminformatics tool PickAxe (34). This tool applies generalized reaction rules to predict potential novel reactions that a given set of metabolites (in this case, all JCVI-Syn3 metabolites) may undergo given known spontaneous (7) and enzymatic(35, 36) chemical mechanisms. We started our PickAxe exploration with the 304 metabolites included in the JCVI-Syn3A model and applied the PickAxe algorithm for multiple iterations to allow for the generation of multistep pathways (see methods). We used both spontaneous and enzymatic reaction rules in the PickAxe expansion, enabling prediction of pathways comprised of a mixture of spontaneous and enzymatic reaction steps (as is the case with damage and repair pathways). In the initial iterations of the PickAxe algorithm, we discovered an increasing number of compounds generated that matched our observed metabolites, but after six iterations, these hits tapered off to just one new compound produced that matched an observed metabolite (blue line in Fig. 6). Interestingly, the number of compounds predicted by PickAxe that matched known biochemistry in the ModelSEED database (green line in Fig. 6) followed a similar trend. We halted the PickAxe expansion at this stage given the diminishing returns in useful or recognizable chemistry being generated. Overall, the final chemical network generated by PickAxe included 33,934 compounds, 61,939 reactions, and matched a total of 182 distinct metabolites (including the original 57 matching the JCVI-Syn3 model) and 1090 ModelSEED compounds (see Supplemental data S4C-D).

**Figure 6.**
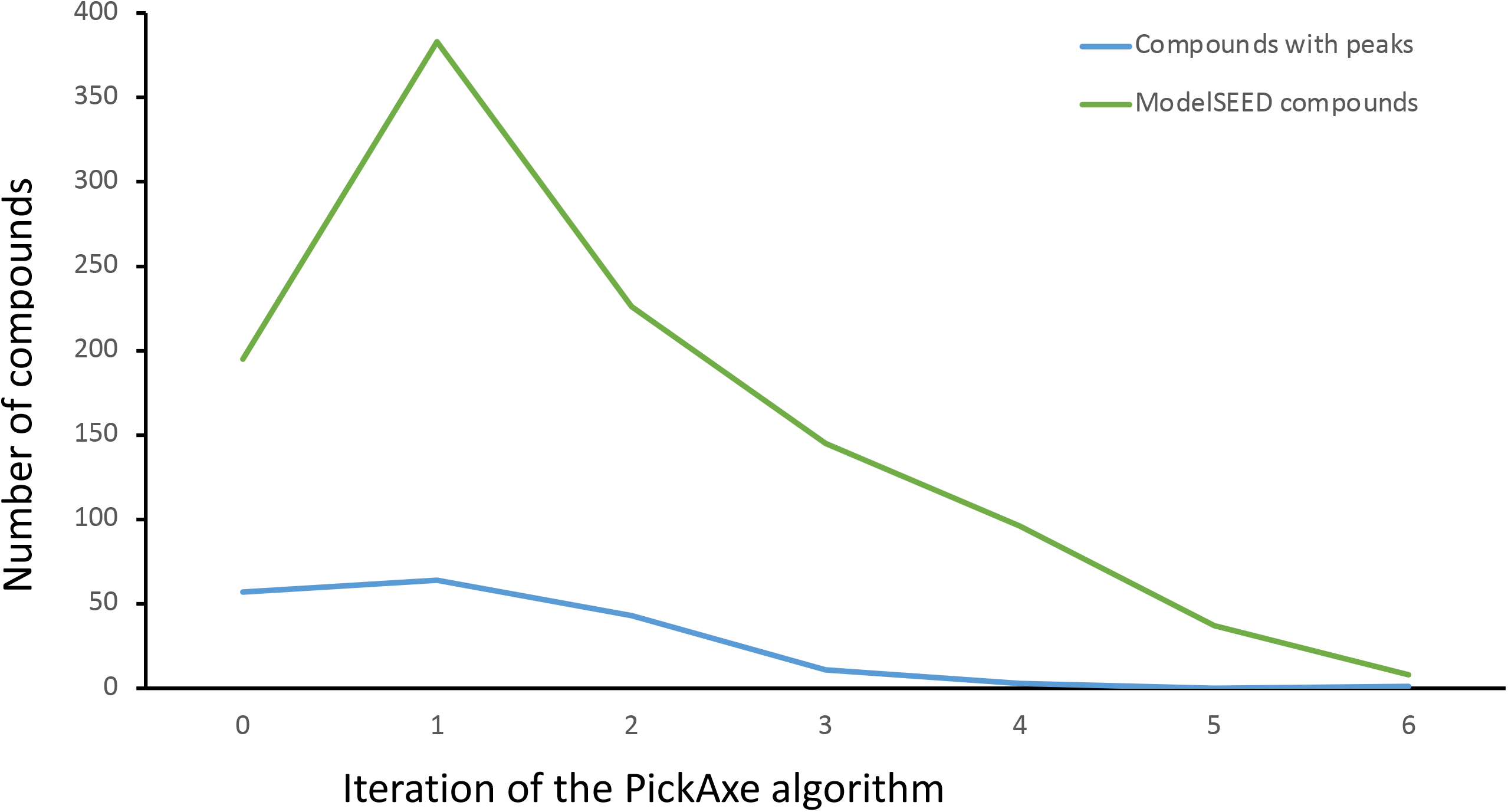
Generation of compounds matching observed peaks (blue line) or ModelSEED compounds (green line) with each iteration of the PickAxe algorithm. Note, multiple compounds can match the same peak if they are stereo isomers.

The network generated by PickAxe represents a pool of hypothetical chemistry possible given our reaction rules and the compounds in the JCVI-Syn3A model. We then used a new flux balance analysis formulation, called metabo-FBA, to select a minimal subset of these reactions that can connect the functioning JCVI-Syn3A model to as many observed metabolites as possible using mass and energy balanced pathways (see Methods). Because we are working with a minimal genome with limited enzymatic diversity and the present study specifically focused on metabolite damage, we favored solutions that involved as many reactions generated by spontaneous reaction rules as possible. Using this approach, we were able to produce a predicted flux profile that succeeded in simultaneously pushing flux through reactions involving compounds that matched 182 distinct observed metabolites (see solution depicted in Fig. 7 and data in Supplemental data S4A and E). This solution included 145 (58%) of the 252 reactions in the JCVI-Syn3 model (purple reactions in Fig. 7), 129 additional ModelSEED reactions (primarily predicted enzymatic reactions; green reactions in Fig. 7), 84 novel enzymatic reactions (blue reactions in Fig. 7), and 74 novel spontaneous reactions (red reactions in Fig. 7) (data in Supplemental data S4A). The fixed image of our flux solution depicted in Fig. 7 is of limited value for permitting a detailed exploration of the fluxes, so we are also including all data files and instructions needed to replicate this view in a fully functioning dynamic Escher map (see Supplemental data S5). Also, the fully expanded version of the JCVI-Syn3A model used to generate this flux solution is provided in SBML and JSON format in Supplemental data S5.

**Figure 7.**
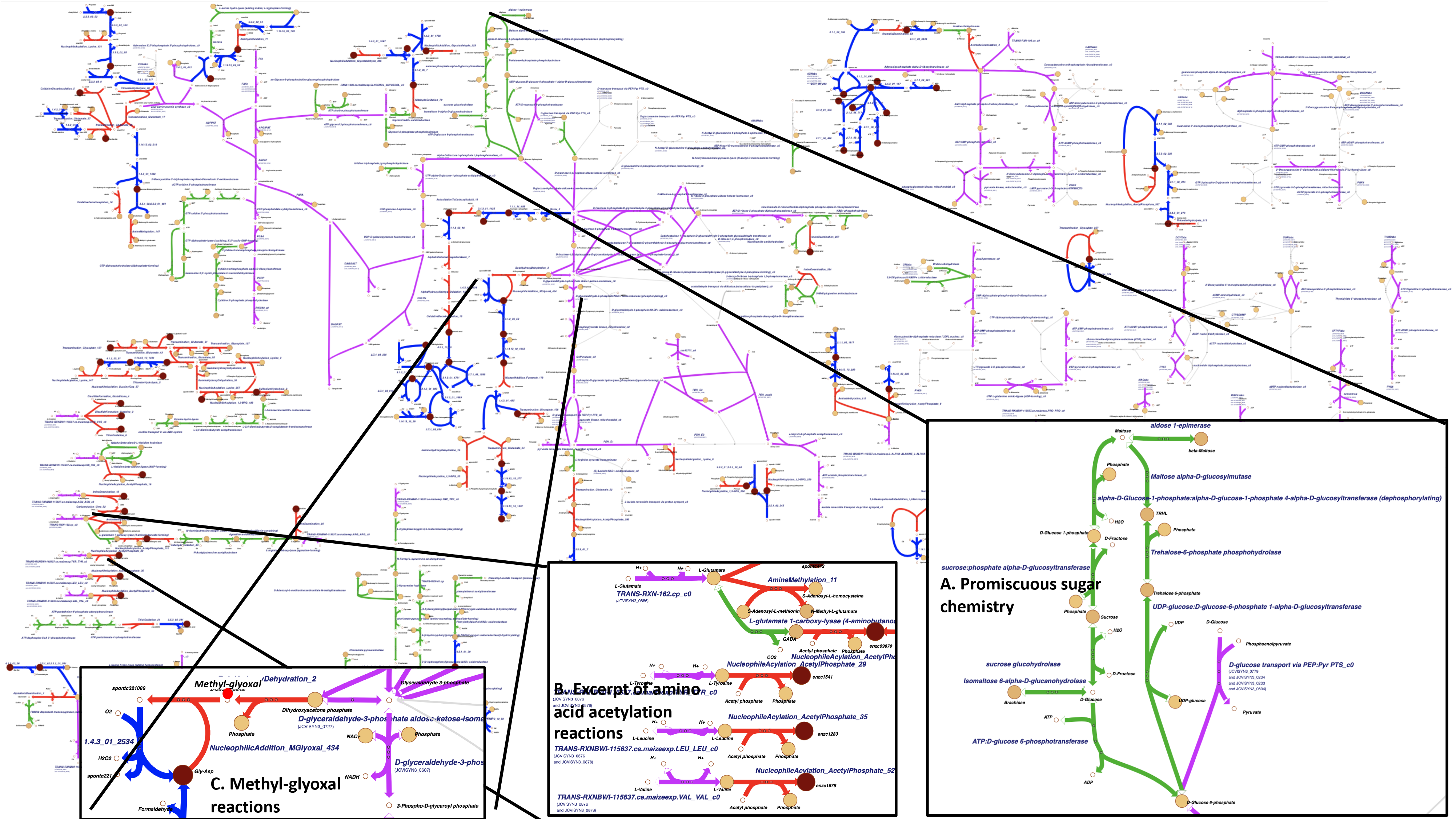
Map of predicted extensions to the JCVI-syn3 model to push flux through as many observed peaks as possible. Grey reactions are inactive model reactions; purple reactions are active model reactions; green indicates active ModelSEED reactions; red indicates active predicted damage reactions; and blue indicates active predicted enzymatic reactions. All active predicted spontaneous reactions and nearly all active model reactions are shown on the map; some ModelSEED and predicted enzymatic reactions are excluded. The insets highlight examples of promiscuous carbohydrate chemistry (A), prevalence of amino acid acetylation (B), and spontaneous reactions mediated by the damage-causing metabolite methylglyoxal.

This flux solution represents only one of many possible solutions to explain the observed metabolomics data based on known and novel biochemical reactions. It is unlikely that this solution is completely correct, but the true solution must make use of similar chemistry, start with the same initial high-confidence JCVI-Syn3A compounds, and produce the same observed metabolic intermediates, meaning the true solution cannot depart too significantly from our selected solution. Thus, while we cannot define exact mechanisms for producing observed metabolites from this analysis, we can observe significant chemical trends that reveal insights into areas of limited understanding of metabolic chemistry and the role of spontaneous reactions in that chemistry.

Looking at the map broadly (Fig. 7), it is immediately apparent that there are hotspots of intense chemical expansion (adenine, cytosine, sugars, pyruvate, amino acids, central carbon trunk reactions, CoA) and other regions with little or no expansion (deoxynucleotides, guanine, thymidine, THF, riboflavin, NAD). This likely has to do with the concentration and reactivity of the associated compounds. For example, deoxynucleotides lack a chemically active hydroxyl group present in standard nucleotides. Many of the intensely branching compounds represent high concentration metabolic starting points (e.g., sugars), end points (amino acids), and high flux intermediates (e.g., pyruvate). Given their higher concentrations, it is more likely that metabolomics will detect these compounds and their derivatives, and that these compounds will undergo additional chemistry.

The large number of ModelSEED reactions and the many predicted novel enzymatic reactions proposed by this approach represent previously unannotated but potential promiscuous side activities of existing annotated gene products in JCVI-Syn3A. The extensive metabolomic evidence for the presence of the products of these reactions points strongly to the presence of the reactions themselves. This is best exemplified by the cluster of ModelSEED reactions expanding from the glucose-6-phosphate (g6p) node of the JCVI-Syn3A model (see Fig. 7A). These reactions involve phosphorylation and hydrolysis interconverting many different sugars and polysaccharides, all of which have evidence for existence in our metabolomics data. While the model only contains reactions for glucose as a representative sugar, it is likely that this model and many other similar models are substantially understating the promiscuity of these enzymes.

Also of note is how many of the pathways predicted in JCVI-Syn3A by our metabo-FBA method involve a mixture of database reactions, predicted spontaneous reactions, and novel enzymatic reactions (30/50 total pathways). Any analysis that focused on only one or even two of these three reaction sources would explain a far smaller number of observed metabolites due to holes and dead-ends in the predicted pathways. A complete understanding of metabolism requires all three reaction data sources.

Another significant trend is the large portion of new predicted chemistry surrounding amino acids. A significant number of observed metabolomics peaks relate to amino acid derivatives, including numerous dipeptides and acetylated amino acids (see Fig. 7B). The dipeptides primarily serve as nutrients for the JCVI-syn3 strain, which contains the peptidases needed to degrade these compounds (a large number of the ModelSEED reactions added by our metabo-FBA approach relate to dipeptide transport and degradation). However, the acetylated amino acids are interesting as only 7 out of 10 of these compounds were found in any biochemistry databases, which also lacked spontaneous reactions for producing these compounds. Yet, metabolomics evidence was found for all 10 compounds being present in the JCVI-Syn3A strain. The metabo-FBA approach added 10 predicted spontaneous acetylation reactions, using acetyl-phosphate as a donor, based on PickAxe predictions. This demonstrates how readily acetylation occurs in these systems, either by spontaneous action or by promiscuous enzyme activity, and it highlights the particular vulnerability of amino acids to this acetylation.

In addition, these results further support certain hypotheses made previously about the main metabolic network of JCVI-Syn3A (3) both with regard to acetyl phosphate as well as the enzymes producing/consuming it. The in vivo essentiality of phosphate acetyltransferase (JCVISYN3_0229) and acetate kinase (JCVISYN3_0230) was puzzling, given that the preceding genes in the pathway, the remaining subunits of pyruvate dehydrogenase (JCVISYN3_0227/8), were found to be non-essential *in vivo*. It had been hypothesized that the two former enzymes thus were not essential because acetate fermentation was essential for the cell, but rather because buildups of acetyl-CoA or acetyl phosphate needed to be prevented, with acetyl phosphate a known protein acetylation agent (37).

Firstly, the current results hence support the role of acetyl phosphate as a biologically relevant acetylation agent in JCVI-Syn3A not only for free amino acids but also for proteins, as some of the identified amino acids had side chain acetylations. Secondly, the results also support the hypothesized essential role of acetate kinase as a means of preventing excess buildups of acetyl phosphate. Thirdly, if acetyl phosphate is indeed the acetylation agent at play, then this implies some source for acetyl phosphate/acetyl-CoA. Furthermore, if at least one of the two hypotheses for the essentiality of phosphate acetyltransferase and acetate kinase is correct (the latter now being supported by the current results), then this source would have to be intracellular production of acetyl-CoA rather than conversion of external acetate (as it would be these two enzymes which would then produce acetyl phosphate and acetyl-CoA in the first place). Production of acetyl-CoA in JCVI-Syn3A had not been certain previously, as the first subunit of pyruvate dehydrogenase and the related NADH oxidase had been removed in JCVI-Syn3A and the remaining components tentatively assumed to still be active. Alternatively, oxidation of acetaldehyde to acetyl-CoA had been hypothesized as a possible function for the remaining pyruvate dehydrogenase complex. The current results would suggest at least one of these two hypothesized pathways to indeed produce acetyl-CoA – or there would have to be yet another mechanism.

We are also particularly interested in using these analyses to understand the relative prevalence, and thus importance, of our various proposed mechanisms for spontaneous chemistry. One can examine prevalence in two ways, and our analyses explore both: (1) how ubiquitous are the active sites that can undergo each given class of spontaneous reaction among the metabolites present in JCVI-Syn3A; and (2) how often can each chemistry be observed to happen based on metabolomics data. We can answer the first question by counting how many reactions are generated by each spontaneous reaction operator in our PickAxe expansion of the JCVI-Syn3A metabolites (orange bars in Fig. 8). From this, we find *carbamylation* to the most dominant spontaneous mechanism with nearly 2000 reactions generated. *Benzoquinone addition* reactions are also very prevalent with almost 1500 reactions generated. However, just because chemistry occurs on a common active site does not mean it will be readily observed. Products from the most visible chemistry are likely to accumulate and thus be observed in metabolomics data. Thus, the reactions selected for addition by our metabo-FBA method represent the most visible in terms of producing significant amounts of observable products. Counting the reactions selected by metabo-FBA for each of our spontaneous reaction mechanisms (gray bars in Fig.8) reveals acylation reactions as being the most common by far, with transamination also being quite common. These results do not necessarily mean that all the other chemistry predicted by our PickAxe analysis outside of those reactions involving or leading to observed metabolites is not happening. Much of this chemistry may still be hidden from analysis if: (1) it only involves compounds with concentrations below the threshold of detection; (2) it involves metabolites that are not easily observed (e.g. highly unstable or volatile compounds); (3) it involves metabolites that are not easily identified (e.g. new compounds with unknown fragmentation patterns and no available standards). Thus, this difference between predicted chemistry and observed chemistry could provide interesting targets for improving methods for observing and identifying metabolites. Additionally, it is important to recognize that many PickAxe reaction types involve co-substrates that are not present in the JCVI-Syn3A strain (e.g. benzoquinone or carbamoyl transferase reactions), and thus it is expected that the products from these reaction types will not appear in JCVI-Syn3A.

**Figure 8.**
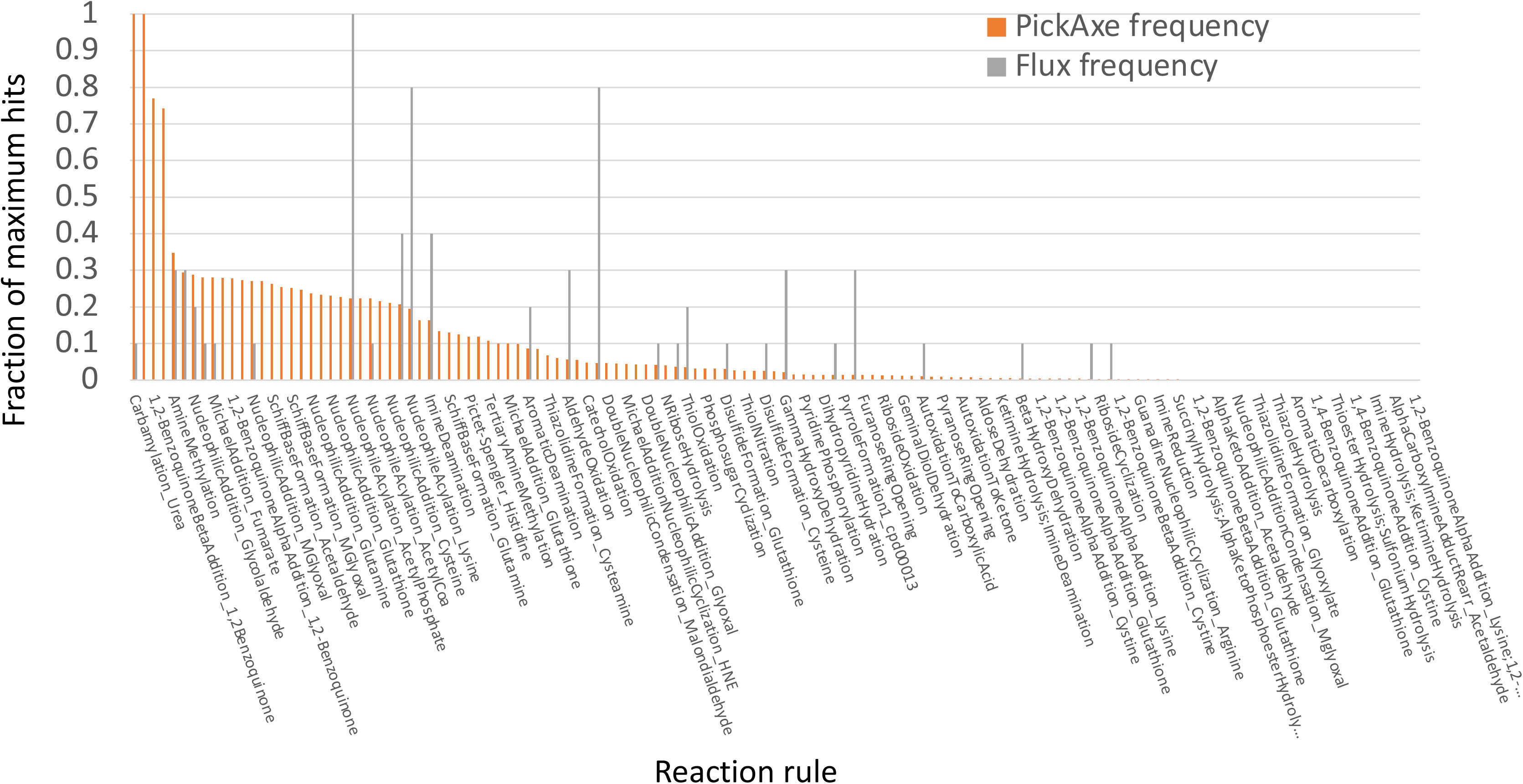
Distribution of predicted and metabolomics-associated reactions generated by spontaneous reaction rules. The orange bars in the chart show the relative number of reactions generated by each spontaneous reaction rule in the full PickAxe expansion of the JCVI-syn3 model. The gray bars show the relative number reactions generated by the same reaction rules that were actually used to generate observed metabolites in the JCVI-syn3 strain by the metabo-FBA analysis. To permit side-by-side comparison, the reaction counts are normalized by the number of reactions associated with the most prevalent reaction rule in each set. The differences in the distributions highlight how the most promiscuous reaction rules (highest orange bars) are not necessarily the most impactful on cell chemistry (highest gray bars).

An example of an important intermediate metabolite that arises from and participates in spontaneous damage reactions but could not be observed using current metabolomics methods was methylglyoxal (see Fig. 7C). While methylglyoxal was not among the observed metabolites due to small size and volatility, metabo-FBA added reactions involving this compound because it leads to numerous downstream potential damage and repair reactions. A more detailed discussion of methylglyoxal follows.

### Exploring possible mechanisms for JCVI-Syn3A to cope with methylglyoxal stress

Methylglyoxal is necessarily generated from the triose phosphates formed by JCVI-Syn3A metabolism (38) but the classical glyoxalase system comprising the glutathione-dependent GloA and GloB enzymes (39) is absent. Likewise, enzymes with minor methylglyoxal-detoxifying activities, such as aldose reductases and keto-aldehyde reductases (40–42) are not encoded in the JCVI-Syn3A genome. The only candidate enzyme that we identified as potentially able to counter methylglyoxal-induced damage is JCVISYN3A_0400, which encodes a homolog of DJ-1. The DJ-1 superfamily has several functionally distinct clades, and phylogenetic analysis places JCVISYN3A_0400 in the YajL/DJ-1 clade (Fig. S6). The clades of the DJ-1 superfamily are not isofunctional and four subfamilies are found in *E. coli* alone (encoded by the *hch*A, *yaj*L, *yhb*O and *elb*B genes).

Although the biochemical functions of many DJ-1 superfamily members remain uncertain, the functionally characterized DJ-1 superfamily proteins are involved in stress response and detoxification mechanisms (43). Some are thought to be deglycases (44), glyoxalases (45) and/or aldehyde-adduct hydrolases (46). We were not able to reproduce the previously reported glyoxal and/or methylglyoxal sensitivities of the Δ*yaj*L/Δ*hch*A *E. coli* K-12 BW25113 strain (44), but we did observe a defect both in its growth rate and yield (Fig. 9A and Fig. S7A). Expression of the *E. coli yajL* or JCVISYN3A_0400 genes *in trans* complemented this growth phenotype (Fig. 9A and Fig. S7A) suggesting JCVISYN3A_0400 was indeed in the same DJ-1 subgroup as YajL.

**Figure 9.**
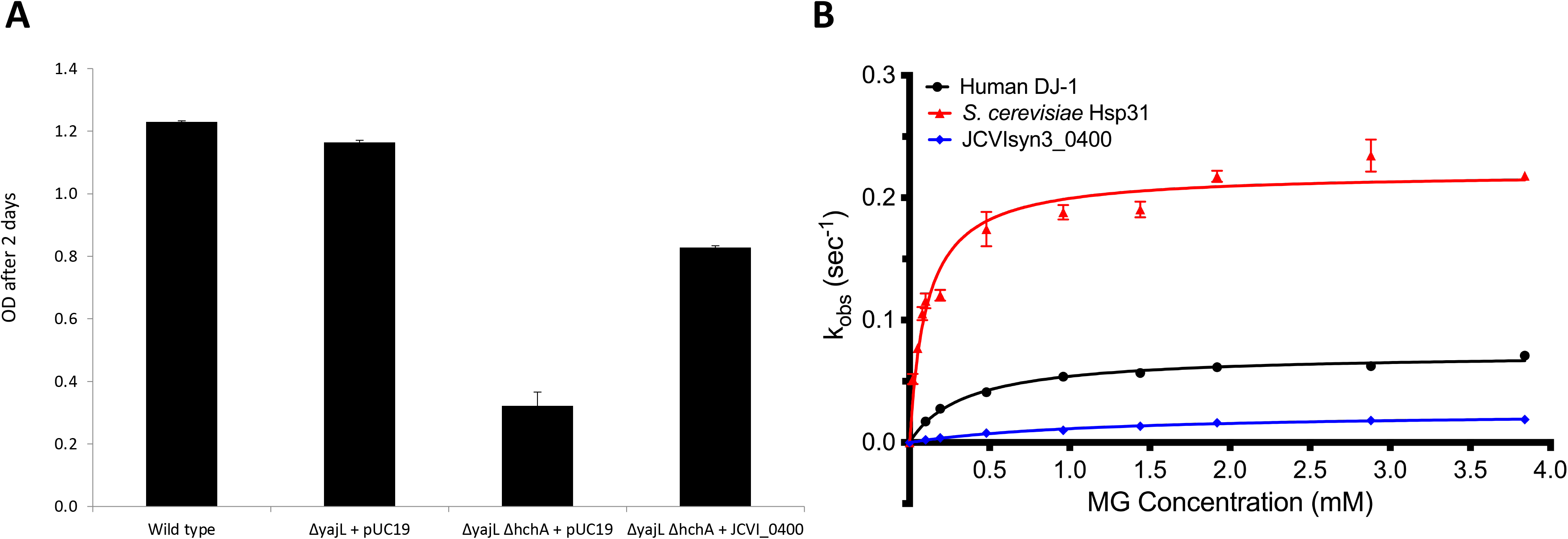
Characterization of JCVI_0400. **(A)** Growth of WT, Δ*yaj*L, Δ*yaj*L Δ*hch*A, Δ*yaj*L Δ*hch*A with *hchA* in trans and Δ*yaj*L Δ*hch*A with JCVI_0400 in trans. pUC19 was used as empty vector. Each strain was tested in 5 replicates Plates were incubated 2 days at 37°C in LB with agitation in a Bioscreen C device. **(B)** Methylglyoxalase activity of JCV_0400 (MP DJ-1) compared to human DJ-1 (HsDj1) and yeast Hsp31. Conversion of methylglyoxal to L-lactate was measured in a coupled assay with L-lactate oxidase and Amplex red. JCVIsyn3_0400 is a weak methylglyoxalase. Hsp31 produces racemic lactate, so the measured k_obs_ is ∼1/2 the true rate and is higher than either close DJ-1 homolog.

To test the hypothesis that JCVISYN3A_0400 is involved in methylglyoxal detoxification we expressed and purified the recombinant protein and measured its glyoxalase activity *in vitro*. As shown Fig. 9B the JCVISYN3A_0400 protein has very low but measurable methylglyoxalase activity (*k*_cat_=0.025±0.002 sec^-1^, *K*_M_=1.23±0.3 mM), far lower than obtained for the positive control proteins *S. cerevisiae* Hsp31 (*k*_cat_=0.220±0.005 sec^-1^, *K*_M_=0.11±0.01 mM) and somewhat lower than human DJ-1 (*k*_cat_=0.073±0.002 sec^-1^, *K*_M_=0.34 ±0.03 mM). The ∼20 M^-1^ sec^-1^ *k*_cat_/*K*_M_ value for JCVISYN3A_0400 is five to six orders of magnitude lower than that of glyoxalase I, the dedicated glutathione-dependent glyoxalase in most organisms (47). Even compared to other DJ-1 superfamily glyoxalases that have relatively low catalytic efficiency, JCVISYN3A_0400 is a poor enzyme. The lactate oxidase-coupled assay used here is specific to L-lactate, which should detect all the lactate produced by JCVISYN3A_0400, as previous reports indicate that DJ-1 clade enzymes produce only L-lactate (48). However, the more proficient Hsp31 glyoxalases produce racemic (D/L)-lactate (48), and thus the rate measured in this assay for *S. cerevisiae* Hsp31 is probably about half the true rate.

Because DJ-1 superfamily members have been reported to be generalist deglycases (49), we tested the deglycase activity of JCVISYN3A_0400 against the methylglyoxal-CoA hemithioacetal (Fig. S7B). CoA was used as the thiol because the absence of glutathione biosynthetic enzymes in JCVI-Syn3A means that CoA may be the principal small molecule thiol in the cell (see above). JCVISYN3A_0400 had no detectable deglycase activity against methylglyoxal-CoA hemithioacetal, while human DJ-1 had a low activity (*k*_cat_=0.021±0.003sec^-1^, *K*_M_=0.39±0.18 mM). Therefore, JCVISYN3A_0400 appears unlikely to efficiently detoxify methylglyoxal via either glyoxalase or deglycase activities if the *in vitro* rates are similar to the *in vivo* activity of the protein. It is possible that JCVISYN3A_0400 and other DJ-1-type glutathione-independent methylglyoxalases may have some unidentified positive effector *in vivo* that could enhance their cellular activity, although there is currently no direct experimental evidence for this. In summary, while results suggest that JCVISYN3A_0400 and YajL are iso-functional, the molecular function of these proteins remain mysterious.

## Conclusion

Metabolite damage arising from side-reactions of enzymes and spontaneous chemistry has often been ignored or seen as a minor metabolic inconvenience that does not warrant investment in enzymes to prevent or repair it (6). Biochemical, genetic, and engineering evidence accumulating over the past decade has been changing this view (6, 7, 10, 12, 50, 51). The biochemical and genetic results we present here constitute particularly persuasive additional evidence by demonstrating that stripping a genome down to its barest essentials leaves metabolite damage-control systems in place. Furthermore, our metabolomic and cheminformatic results point to the existence of a network of metabolite damage and damage-control reactions that extends far beyond the corners of it characterized so far. In sum, there can be little room left to doubt that damage itself and the systems that counter it are mainstream metabolic processes.

## Methods

### Bioinformatics

The BLAST tools (52) and CDD resources at NCBI (http://www.ncbi.nlm.nih.gov/) (53) were routinely used. Sequences were aligned using Clustal Omega (54) or Multialin (55). Phylogenetic distribution was analyzed in the SEED database (56). Results are available in the “YqeK” subsystem on the PubSEED server (http://pubseed.theseed.org//SubsysEditor.cgi?page=ShowSpreadsheet&subsystem=NadD-YqeK_fusion_display). Physical clustering was analyzed with the SEED subsystem coloring tool or the SeedViewer Compare Regions tool (56) and the clustering figure was generated with GeneGraphics (57). Phylogenetic trees were constructed with Mega 6 (58). Student’s t-test calculations were performed using the VassarStats web-tools (http://vassarstats.net).

### Prediction of novel potential chemistry using PickAxe

Expanded chemistry was generated using the PickAxe app in KBase, as shown in this narrative: https://narrative.kbase.us/narrative/29280. This app uses the open source RDKit package to apply sets of SMARTS-based chemical reaction rules, derived from previously published chemical damage (7) and enzyme promiscuity (34) studies, to an input set of compounds to produce all possible reactions and products that might arise from that chemistry. This analysis can be run iteratively through repeated application of the reaction rules to all new products that arise from previous generations. We applied the PickAxe approach for six iterations, retaining all compounds that matched the JCVI-Syn3A model, the ModelSEED database (31), or an observed metabolite.

### Metabo-flux balance analysis to predict minimal reactions to reach observed metabolites

In metabo-flux balance analysis (metabo-FBA), constraints are added to the standard FBA formulation to force flux through one or more reactions involving an observed metabolite. In this formulation, a variable is added for each observed peak (p_i_) and a variable is added for each metabolite that has been mapped to the peak (because peaks lack stereochemistry, they may be mapped to multiple possible stereoisomers). Next, a constraint is added stating that a peak cannot be active unless one or more of its associated metabolites is active (where *λ*_*i*,*j*_ is a mapping variable equal to 1 if metabolite *j* is mapped to peak *i* and zero otherwise):

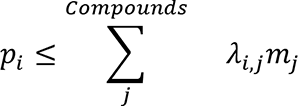

A constraint is also added stating that no metabolite can be active unless at least one reaction in which the metabolite is involved is carrying flux (where *γ*_*j*,*k*_ is a mapping variable equal to 1 if metabolite *j* is involved in reaction *k* and zero otherwise):

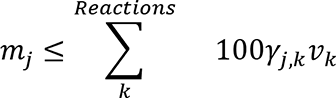

To maximize active metabolites, the objective of the problem is then set to maximize the sum of all *p_i_*. While *p_i_* and *m_j_* can be specified as binary variables, it works equally well and is less computationally expensive to use continuous variables bounded between 0 and 0.1. To avoid the trivial solution of activating metabolites by pushing flux through both directions of reversible reactions or around mass balanced flux loops, it is essential to also employ thermodynamics constraints in some form in this formulation (59).

### Synthesis of NaAD^+^ analogs and nicotinic acid riboside (NaR)

NaMN (0.5 mM), 4 mM MgCl_2_, 5 units/ml yeast inorganic pyrophosphatase, 1 mg/ml BSA and 2 mM (d)NTP (dATP, CTP, GTP or UTP) were incubated with 150 μg NadD-YqeKH230A enzyme overnight at 37° C in 20 mM HEPES-KOH, pH 7.2, 100 mM NaCl, 0.2 mM DTT, 1% glycerol. Assays were deproteinized using Amicon 10K cutoff centrifugal filters, concentrated *in vacuo* and purified by HPLC (Waters 2695 Separation module and Waters 2998 PDA detector) using a C18 column (Thermo Scientific Hypersil GOLD C18 5 μm, 250×4.6 mm) with a column guard with 20 mM ammonium bicarbonate / acetic acid, pH 6.0. Purified NaAD^+^ analogs were lyophilized and resuspended in 10 mM HCl, pH 2.0.

To synthesize a NaR standard, NaMN (10 mM) was dephosphorylated with 20 units CIP overnight at 37° C. The mixture was deproteinized using Amicon 10K cutoff centrifugal filters and used as a standard as is. The following extinction coefficients were used to quantify NaAD^+^ and its analogs: NaAD^+^ 19.4 ×10^-3^ M^-1^, dNaAD^+^ 19.4 ×10^-3^ M^-1^, NaCD^+^ 11.9 ×10^-3^ M^-1^, NaGD^+^ 16.4 ×10^-3^ M^-1^, NaUD^+^ 13.3 × 10^-3^ M^-1^. Extinction coefficients were based on published extinction coefficients of NAD^+^ analogs (60). To adjust for the nicotinic acid moiety, the difference of extinction coefficients of nicotinic acid (4.2 × 10^-3^ M^-1^) and nicotinamide (2.78 × 10^-3^ M^-1^) (61)was added to those of the published NAD^+^ analogs.

### Media, strains, and genetic manipulations

All strains, plasmids and oligonucleotides used in this study are listed in Table S4 and Table S5. Bacterial growth media were solidified with 15 g/l agar (BD Diagnostics Systems) for the preparation of plates. *E. coli* were routinely grown on LB medium (BD Diagnostics Systems) at 37 °C unless otherwise stated. Transformations were performed following standard procedures (62). IPTG (100 μM), Ampicillin (Amp, 100 μg/ml), Kanamycin (Km, 50 μg/ml), l-Arabinose (Ara, 0.02–0.2%), Chloramphenicol (Cm, 25 µg/ml) and Rifampicin (Rif, 25 µg/ml) were used when appropriate. Bacterial M9 minimal medium (62), 0.4% (w/v) glucose was used either with NH_4_Cl (20 mM) or glycine (50 mM) as the nitrogen source. P1 transduction was performed following the classical methods (63). The Kan^R^ marker was eliminated from the BW2113 Δ*yaj*L::Kan^R^ strain by the procedure described by Cherepanov and Wackernagel (64). Transductants from BW2113 Δ*hch*A::Kan^R^ to BW2113Δ*yaj*L were checked by PCR for transduction of the Δ*hch*A::Kan^R^ allele into the recipient strains using primer pairs [DH492/493 (ext); DH494/495 (int) and DH480/481 (ext); DH482/483 (int)] respectively.

JCVI-syn3A is a near minimal bacterial cell first reported by Breuer et al. (3)that contains a subset of the genes in *Mycoplasma mycoides* subspecies *capri* strain GM12. Mycoplasmas were grown in SP4 broth (65) that contains 17% KnockOut Serum Replacement™ instead of 17% fetal bovine serum and is referred to as SP4-KO as described in the supplemental Methods.

### Construction and analysis of JCVI_Syn3A hydrolase gene deletion mutants

Construction of gene knockout mutants in JCVI-Syn3A was a multistep process, and two different protocols were used. Protocol I entailed CRISPR/Cas9 mediated removal of an individual target gene from a JCVI-Syn3A genome cloned as a yeast centromeric plasmid (YCp) in yeast strain VL648NCAS9_Syn3A that carries the *cas9* gene in the yeast genome and expresses Cas9 constitutively (2, 66, 67). Following the CRISPR/Cas9/homologous recombination with a donor DNA to re-circularize the JCVI-syn3A YCp, each mutated genome was transplanted by standard procedures (68, 69) in order to produce JCVI-syn3A bacteria lacking the respective target gene. Protocol I successfully removed gene JCVISYN3A_0728 but did not yield bacterial deletion mutants for any of the other four hydrolase genes. Protocol II, which is more complicated, was then used to make deletion mutants for the other hydrolase genes JCVISYN3A _0066, JCVISYN3A _0077, JCVISYN3A _0710, and. JCVISYN3A _0907. Using the yeast deletion constructs generated in JCVI-syn3A YCp using Protocol I, a second CRISPR/Cas9 was applied to install each respective hydrolase gene behind its native promoter between loxP sites in a non-essential region of the JCVI-syn3A genome. These four YCps were successfully transplanted to generate individual JCVI-syn3A bacteria, each with the respective target gene in essentially the same new location. We then transformed these bacterial relocation mutants, harboring the target gene between loxP sites, with a plasmid containing i) a Cre recombinase gene under transcriptional control of a mycoplasma promoter and (ii) a puromycin resistance gene. With this plasmid positioned between the same two loxP sites in the relocation mutants, cells that were plated on Sp4 growth media containing puromycin, efficiently exchanged the puromycin resistance cassette for genes JCVISYN3A _0066, JCVISYN3A _007, and JCVISYN3A _0907, yielding the desired bacterial deletion mutants. We did not obtain a JCVISYN3A _0710 deletion mutant. These protocols are described in detail in the **Supplemental data S2** file.

### Inactivation of the hydrolase encoding domain of gene JCVISYN3A _0380 by converting the His codon at position 230 to an Ala (His230Ala)

This was done using CRISPR/Cas9 as described previously to cut a yeast clone of the JCVI-syn3A YCp in gene JCVISYN3A _0379. Next, while the CRISPR cut JCVI-syn3A YCp was still in yeast, a 1727 bp DNA molecule made using a multistage PCR that encoded gene MMSYN1_0380 with the desired His to Ala mutation at codon 230 and flanked by sections of genes JCVISYN3A _0379 and JCVISYN3A _0381 was recombined into the linear JCVI-syn3A YCp. The resulting mutated plasmid was installed in a *Mycoplasma capricolum* cell using genome transplantation to create the JCVI-syn3A with the mutated JCVISYN3A _0380. The process is described in detail in the Supplemental data S2 file.

### Plasmid constructions for expression JCVI-syn3A genes in *E. coli*

The sequences encoding all the JCVI-syn3A genes characterized in this study were codon-optimized by the supplier (GenScript, Piscataway, NJ) for expression in *E. coli*. They were synthesized with added restriction sites at the 5′ and 3′ ends and cloned in different vectors: pUC19 for JCVISYN3A_0400, JCVISYN3A_0443 and JCVISYN3A_0887 at SphI and NcoI sites; pUC57 for JCVISYN3A_0380 at NdeI and XhoI sites, or pET28a (JCVISYN3A_0728, JCVISYN3A_0066, JCVISYN3A_0077, JCVISYN3A_907) at XbaI and XhoI sites which added a C-terminal His-tag. The nucleotide sequences of all synthesized genes are given in Supplemental Methods and the corresponding plasmids listed in Supplemental Table S4. The sequences of the oligonucleotide primers used to subclone certain synthesized genes from pUC19/pUC57 to other vectors are given Supplemental Table S5. JCVISYN3A_0887 was cloned in the NcoI and BlpI sites of pET28 which added a C-terminal His tag after PCR amplification from pUC19-887 using the DH526 and DH527 primers. JCVISYN3A_0380 was cloned in the NcoI and PstI sites of pBAD24 after PCR amplification from pUC57-380 using the DH540 and DH541 primers. The same restriction sites were used to clone in pBAD24 the fragment encoding only the NadD_M_ domain (amplified by PCR from pUC57-380 using the DH540/DH584 primer pair) and the YqeK_M_ domain (amplified by PCR from pUC57-380 using the DH585/DH541 primer pair). Design of the primers to separate the domains was based on multiple alignment shown Fig. S4B. The JCVISYN3A_0380 was sub-cloned from pUC57-380 after digestion with NdeI and XhoI restriction sites and ligated into the matching restriction sites of pET28a, which adds an N-terminal His tag.

*Bs*NadD_Bs_ was PCR amplified with Phusion High-Fidelity DNA polymerase (New England BioLabs) from *Bacillus subtilis* 168 genomic DNA using primers BsNadD_Fw and BsNadD_Rv-XhoI digested with NdeI/XhoI and ligated into the matching sites of pET28b (Novagen), which added a N-terminal His_6_-tag. The syn3.0 NadD-YqeK gene was synthesized by Genscript. The YqeK domain was cloned into NdeI und XhoI restriction sites of pET28 after PCR amplification using primers synYqeK-Fw_NdeI and synYqeK-Rv_XhoISTOP, which adds a N-terminal His-Tag. The YqeK domain was cloned into NdeI und XhoI restriction sites of pET28 after PCR amplification using primers synYqeK-Fw_NdeI and synYqeK-Rv_XhoISTOP, which adds a N-terminal His-Tag. The JCVIJCVISYN3A_0380 H230A mutant was made by site directed mutagenesis, using primers H230AFor/H230ARev to change the CAC codon (His) to GCC (Ala), using Phusion polymerase. JCVISYN3A_0400 was cloned between the NdeI and BamHI sites of bacterial expression vector pET15b after PCR amplification from pUC19-400 using primers DH611 and DH612. All constructs were verified by Sanger sequencing.

### Mutation frequency assays for *E. coli* derivatives

Overnight cultures in LB with added antibiotics and arabinose (0.02%) were diluted 100-fold in the same conditions and grown for another 24 h before dilutions were plated on LB and LB rifampicin (25 μg/ml) to calculate a mutation ratio (Number of colonies on Rif x dilution factor) / (Number of colonies on LB x dilution factor).

### Protein expression and purification and enzyme assays

All characterized JCVI-syn3A encoded proteins were expressed as His-tagged variants in *E. coli* and purified using Ni^2+^-NTA columns as described in Supplemental Methods. In vitro activity assays for CoA disulfide reductase, for phosphatase with a range of substrates, NadD, glyoxalase, and deglycase are described in detail in Supplemental Methods.

## Acknowledgements

This work was funded by the National Science Foundation (Grants MCB 1611846 to OF, MCB-1611952 to CH and MCB-1611711 to ADH and V dC-L, MCB 1840301, MCB 1840320 and MCB 1818344 subcontracts to J.I.G.) and by the J. Craig Venter Institute.

