## Supplementary material for "Metabolite Damage and Damage-Control in a Minimal Genome": Sup data 2

**Supplemental data S2**

**Culture, growth rate measurements and sampling of JCVI-syn3A and derivative cells.**

JCVI-syn3A and derivative mutant strains used in this study were grown as described in detail elsewhere (1, 2). Briefly, static planktonic liquid cultures were maintained without antibiotic selection by serial passage at 37°C in SP4 liquid medium (3) containing either the standard addition of 17% (v/v) fetal bovine serum (FBS) or, for cells used in metabolomic analyses, the alternative addition of 17% (v/v) liquid serum substitute KnockOut™ (Life Technologies) (SP4-KO).

Growth rates and the stage of cultures were measured as described elsewhere (1, 2) by monitoring cell-associated nucleic acid using a fluorescent dye. Briefly, samples of culture obtained at different times were centrifuged through sucrose cushions to pellet cells and efficiently remove medium. Cells were then lysed with dilute detergent, and the released dsDNA (and to a lesser extent dsRNA) was measured using the fluorescent stain Quant-iT PicoGreen (Molecular Probes, Eugene, OR). Fluorescence was measured in a 96-well format using a FlexStation 3 fluorimeter (Molecular Devices, San Jose, CA). The net relative fluorescence units (RFU) of cell lysates (after subtracting RFU from a mock medium control lacking cells), were plotted as ln(RFU) vs. time from which the doubling times, $\tau$d were calculated from the slopes of exponential regression curves ($k$) as $\tau d=\ln 2/k$. Rates were measured from log–linear portions of the growth curve. To avoid variables such as batch differences among medium preparations and temperature fluctuations, mutants within a group (i.e. the mutants with gene deletions vs. mutants with single nucleotide substitutions) were compared under identical conditions within a single experiment. The high accuracy and reproducibility of rate measurements, as reflected by $R^2$ values of all regression curves, allowed the use of single samples for these comparisons.

Preparation of cells for metabolomics measurements was designed to maintain the physical integrity of JCVI-syn3A and derivative cells during manipulation, minimize the time of preparation, normalize the amounts of sample analyzed, and ensure statistical significance. For each mutant or control strain, cells grown in SP4-KO to log phase were seeded at estimated dilutions into three biological replicate cultures of 8 mL and growth was monitored as described above. At a time determined to be within log phase culture for each biological replicate, all samples within a mutant group were harvested concurrently by placing tubes in wet ice, and final net RFU values were determined for each replicate. Keeping all subsequent steps at 0-4˚C, three identical aliquots from each biological replicate culture were transferred to microfuge tubes to generate technical replicates, adjusting the aliquot volume taken from each biological replicate in order to normalize the cell content among all samples. Tubes were centrifuged at 10,000 x g for 10 min, supernatants were carefully aspirated, and pellets were then rinsed *in situ* with 1.0 mL Tris-saline (TS) buffer (0.15 M NaCl, 20 mM Tris, pH 7.5), which was then thoroughly removed by aspiration. Finally, tubes were plunged into liquid nitrogen to flash freeze cells, pellets were overlaid with 50 µL methanol (Optima LCMS, Fisher), and tubes were sealed and placed on dry ice for transfer to storage at -80˚C. Aliquots of unused SP4-KO medium were frozen at -80˚C to serve as a control to identify the many possible metabolites likely present in the complex SP4 medium required to support JCVI-syn3A growth.

**Construction and analysis of JCVI_Syn3A HAD encoding gene deletion mutants.** Construction of gene knockout mutants in JCVI-Syn3A was a multistep process, and two different protocols were used. The first, initially entailed CRISPR/Cas9 mediated removal of the target gene from a JCVI-Syn3A genome cloned as a yeast centromeric plasmid (JCVI-syn3A YCp) in yeast strain VL648NCAS9_Syn3A that carries the *cas9* gene in the yeast genome and expresses Cas9 constitutively (2, 4).

***Protocol I: CRISPR/Cas9 deletion of target genes***. For this process, we first made synthetic single guide RNAs (sgRNA) by *in vitro* transcription with the T7 RiboMAX™ Express Large Scale RNA Production System (Promega corporation). To construct the *in vitro* transcription template, we produced 119-120 bp PCR amplicons using a three oligonucleotide reaction comprised of an 83 base oligonucleotide encoding a CRISPR/Cas9 guide RNA sequence (sgRNA_Template) that serves as the PCR template (IDT Ultramer), a forward primer comprised of a T7 RNA polymerase promoter at the 5' end, 19-20 bases of gRNA target sequence from the gene to be deleted in the middle, and 22 3' end bases that will anneal to the 5' end of complement of the sgRNA, and a reverse primer (gRNA_R) that is complementary to the 19 bases at the 3' end of the sgRNA. The two shorter oligonucleotides were standard IDT oligos and sgRNA_Template was an IDT Ultramer (Table SA1)(5). We re-suspended the three oligonucleotides in RNase and DNase free water to a final concentration of 10 μM. The 25 μl PCR was comprised of 12.5 μl 2X Q5® DNA polymerase Master Mix (New England Biolabs), 1 μl of each primer (10 μM), 10.3 μl $H_2O$, and 0.2 μl (10 ng) of the 83 base sgRNA template. The PCR began with 2 minutes at 98° C and then had 30 cycles of 98° C for 10 seconds, 55° C for 10 seconds, and 72° C for 10 seconds, followed by 72° C for 5 minutes. We then purified the PCR amplicons using QIAquick columns (Qiagen). We used 8 μl of the PCR amplicons as template in a 20 μl transcription reaction with the T7 RiboMAX™ Express Large Scale RNA Production System according to the manufacturer's instructions. Briefly, we combined 10 μl of RiboMAX™ Express T7 2X Buffer, 158 ng of the previously generated PCR amplicon (in 8 μl $H_2O$), and 2 μl Enzyme Mix T7 Express (Promega). The transcription reaction was incubated at 37 °C for 4 hours or overnight followed by addition of 1 μl of the Turbo DNase provided with the kit and incubation at 37 °C for another 15 min. The reaction volume was adjusted to 750 μl with RNase-free water and 1/10$^{th}$ volume of 3M sodium acetate, followed by phenol-chloroform extraction. RNA was precipitated following this with two volumes of 100% ethanol and washed with 500 μl of 70% ethanol. The pellet was air-dried and then re-suspended in 40 μl RNase-free water.

Next, we transformed yeast cells containing the JCVI-syn3A YCP with single guide RNAs and long donor cassette DNA patches via electroporation. Briefly, 50 ml of yeast culture was inoculated with yeast colony from yeast agar plate and incubated at 30 °C for 24 hours with vigorous shaking. We harvested the culture by centrifugation at 2000 *g* for 3 min. The pellet was washed twice with 50 ml cold, sterile distilled water, and re-suspended in 800 μl of 100 mM lithium acetate solution in 1X TE buffer (pH 8) to make the cells competent for transformation. We added 20 μl of 1 M DTT and incubated the cells at 30 °C for 45 min with gentle shaking at 100 rpm. Cells were washed with 1 ml of cold sterile water, followed by a wash with 1 mL of ice-cold 1 M sorbitol solution, and finally re-suspended in 500 μl of 1 M sorbitol solution. We used 100 μl of cells per electroporation with 1 μg of each single guide RNA, 0.5–1 μg of single donor cassette DNA patches (Supplementary Table SA2; IDT Ultramers) or a pool of donor cassette patches (these seal the Cas9 cleaved YCPs), and 100 ng of a selection plasmid, pYAC_TRP1(4) (makes cells capable of growth on media without tryptophan). We used Gene Pulser®/MicroPulser™ Electroporation Cuvettes, 0.2 cm gap (Bio-Rad) and electroporated cells under the following conditions: 2.5 kV, 200 Ohms, 25 μF. Electroporated cells were transferred into 1 ml of 1:1 mix of 1 M sorbitol/YEPD media, incubated at 30 °C for 2 hours with shaking and plated on YEPD agar selection plates that lacked tryptophan and histidine. Plates were incubated at 30 °C for 3-4 days and several transformants were patched on to selection plates without histidine before screening for the chromosome editing event (Table SA3) and the integrity of the JCVI-syn3A YCp. That screening was done using multiplex PCRs with the QIAGEN multiplex PCR kit according to the manufacturer's instructions (diagnostic PCR primers that amplify 8 regions around the JCVI-syn3A YCp and JCVI-syn3A genome are listed in Supplementary Table SA4). That mutated genome was then tested to determine if it could support a living cell using a genome transplantation process where the mutated YCPs were installed in *Mycoplasma capricolum* recipient cells (6, 7)  using a *tetM* selectable antibiotic resistance marker present in the YCP for the selection of the transplanted cells. Transplants were selected after plating on agar plates containing SP4 glucose broth (3) plus 17% KnockOut serum replacement (Thermofisher) and tetracycline (4 μg/ml). Using Protocol I, we were only able to make the JCVISYN3A_0728 gene knockout. The other hydrolase genes targeted for deletion, JCVISYN3A_0066, JCVISYN3A_0077, JCVISYN3A_0710, and JCVISYN3A_0907 did not yield knockout mutants using Protocol 1. The failure to obtain a JCVISYN3A_0710 deletion mutant was not surprising given that the gene is essential based on transposon bombardment studies; however, JCVISYN3A_0066, JCVISYN3A_0077, and JCVISYN3A_0907 are quasi-essential and we thought that we might obtain mutants (2). We reasoned that because of the low efficiency of the genome transplantation technique (~1 in 100,000 *M. capricolum* cells produce transplants (7) might not have recovered gene knockouts because they are rare, and/or may also grow too slowly for isolation using genome transplantation.

*Protocol II.* Consequently, we employed a second more complicated protocol when the initial method failed that would not depend on obtaining gene knockouts via genome transplantation. In this approach, the gene to be deleted was first removed from its normal position in the JCVI-syn3A YCp as in Protocol I. Then, the gene was reinstalled in same JCVI-syn3A YCp at neutral site in the YCp between two loxP recombination sites using a CRISPR/Cas9 approach as in Protocol I. This YCp that contained the target gene in a new location was then booted up via genome transplantation. Next in the bacteria resulting from the transplantation reaction, we swapped the target gene, which was located between loxP sites for a puromycin antibiotic resistance marker in a highly efficient reaction catalyzed by Cre recombinase.

For the CRISPR/Cas9 stage of this protocol, synthetic sgRNA templates were constructed to produce sgRNAs that would cleave the JCVI-syn3A YCp between the HIS3 yeast selection marker (JCVISYN3A_0918) and the *tetM* bacterial selection marker (JCVISYN3A_0913). As in Protocol I, we did a PCR using the 83 base sgRNA_Template as template and a forward primer, His_gRNA_primer F, and reverse primer, gRNA_R (Supplementary Table SA5). Other than using a different forward primer, conditions for the PCR and *in vitro* transcription of the sgRNA using the T7 RiboMAX™ Express Large Scale RNA Production System were the same as described in Protocol I.

Long donor cassette DNA molecules flanked by loxP sites and containing the target gene, regulatory sequences for transcription and translation, and edges that will hybridize to the JCVI-syn3A YCp/genome sequences flanking the loxP sites were made using two rounds of PCR (primers listed in Supplementary Table S6). These were done for target genes JCVISYN3A_0066, JCVISYN3A_0077, JCVISYN3A_0710, and JCVISYN3A_0907. Using construction of a long donor cassette DNA for gene JCVISYN3A_0066 an example, the procedure was as follows: in the first round of PCR lox66 F1 and lox66 R1 primers, and JCVI-syn1.0 genomic DNA as template were used to amplify gene JCVISYN3A_0066 (25 μl PCR comprised of 12.5 μl 2X PrimeSTAR Max Premix (Takara), 1 μl of each primer (0.2 μM final), 10.3 μl $H_2O$, and 0.2 μl (200 ng) JCVI-syn3A template). The 3' 21 bases of the primers hybridize outside the target genes (for other target genes the hybridization region may be longer). The middle 34 bases of the primers are the loxP sites, and the 5' 24 bases are sequences from the yeast *met14p* gene. The PCR began with 2 minutes at 98 °C and then had 30 cycles of 98 °C for

10 seconds, 55 °C for 10 seconds, and 72 °C for 10 seconds, followed by 72 °C for 5 minutes. The PCR amplicon was purified using a Zymo column (Zymo Research). That purified amplicon was then used as template in the second round of PCR. For all target genes the second PCR round used His land F and ARS land R as primers (Supplementary Table SA6). The PCR products of the second round PCR were purified using Zymo column. In the second round, the 3' 23 bases of the primers hybridized to the yeast *met14p* sequences at the termini of the first PCR amplicon the 5' 55 bases identical to sequences between the JCVI-syn3A genome *his3* and *tetM* genes on either side of the CRISPR/Cas9 cut (Supplementary Table SA7).

Next, we transformed yeast cells containing the JCVI-syn3A YCPs with one these genes JCVISYN3A_0066, or JCVISYN3A_0077, or JCVISYN3A_0710, or JCVISYN3A_0907 deleted with single guide RNAs and long donor cassette molecules containing loxP sites flanking the target genes via electroporation. This was done as in Protocol 1, but we used pRS316 (ATCC® 77145™), for its *ura3* selective marker. We plated the CRISPR/Cas9 reaction mix on CM agar plates lacking uracil and histidine (CM Glucose media, Dry, w/o Histidine and Uracil; Teknova) with 2% agar to make plate. Following genome transplantation using the modified yeast genomes, we did colony PCRs on the resulting mycoplasma colonies as in Protocol 1 using the primer sets listed in Supplementary Table SA7. Properly mutated JCVI-syn3A YCps where the target genes were located between loxP sites were used in genome transplantation reactions as described previously. Then transplants were transformed with the plasmid Pmod2*lox*puro*lox*-cre-sp (GenBank accession number MN982903) that encodes a Cre recombinase gene and also a puromycin resistance protein gene located between the same two loxP sites that flank the target gene. Expression of the Cre recombinase resulted in Cre/*lox*-based recombination mediated cassette exchange of the puromycin resistance marker for the target gene. If the target gene was non-essential, then JCVI-syn3A bacteria lacking the target gene are obtained. The process was very efficient. Thousands of colonies were obtained even if only 100 ng of Pmod2*lox*puro*lox*-cre-sp plasmid was used for a transformation. Using this protocol, we obtained JCVI-syn3A strains lacking genes JCVISYN3A_066, JCVISYN3A_077, and JCVISYN3A_0907. Not unexpectedly given gene JCVISYN3A_0710 was not disrupted in transposon bombardment studies, we were unable to obtain bene JCVISYN3A_0710 knockouts. This confirmed the essentiality of that hydrolase. The transformation of *M. mycoides* was described previously (2).

Basically, the *M. mycoides* cells were grown in 4 ml of Sp4 media to reach pH 6.5-7.0. The cultures were centrifuged for 15 min at 4700 rpm at 10 °C in 50 ml tubes. We resuspended cell pellets in 3 ml of sucrose 0.5 M, Tris 10 mM pH 6.5 (buffer S/T). The resuspended cells were centrifuged for 15 min at 4700 rpm at 10 ºC. The supernatants were removed, and we resuspended pellets in 250 µl of 0.1 M $CaCl_2$ and incubated them for 30 min in ice. We then added 200 ng of plasmid to each tube of cells and mixed very gently. Then to each sample, we added 2 ml of 70% polyethylene glycol (PEG) 6000 (Sigma) dissolved in S/T buffer and mixed well during 2 min of incubation. Immediately thereafter, 20 ml S/T buffer was added and mixed well. The tube was centrifuged for 15 min at 10,000 x g at 8°C. Discarded the supernatants and then we put the tubes upside down on Kimwipes paper, so the PEG flowed out of the tubes. The cells were resuspended well in 1 ml of warm Sp4 media. These cells were incubated for 2 hours at 37 °C and then plated in Sp4 agar with 3 ug/ml of puromycin (Sigma). Colonies appeared after 3-4 days at 37 °C.

**Inactivation of the hydrolase encoding domain of gene JCVISYN3A_0380 by converting the His codon at position 230 to an Ala.**

Mutation of the JCVISYN3A_0380 gene in the hydrolase domain by converting codon 230 from a His to an Ala (CAC to GCA; hereafter called His230Ala) was done in a JCVI-syn3A YCp using two rounds of CRISPR/Cas9 and yeast homologous recombination with a donor DNA, followed by genome transplantation. To ensure that these manipulations did not affect the final construct a control mutation, termed His230His, was made using the same procedure to change the CAC codon to the synonymous CAT codon.

In the first CRISPR/Cas9 step, the molecule to be mutated was cleaved and the donor DNA oligonucleotide 380_40bp_161L recombined with the cut JCVI-syn3A YCp removing parts of genes JCVISYN3A_0379 and JCVISYN3A_0381 and all of gene JCVISYN3A_0380. To construct a template for this CRISPR/Cas9 step sgRNA, PCR was performed using 380gRNA_F5 and gRNA_R as primers and the 83 base sgRNA_Template Ultramer as template to cut the genome in gene JCVISYN3A_0379 (Supplementary Table S8). The PCR conditions and protocol for *in vitro* transcribing the sgRNA are as described previously in Protocol I to make sgRNAs. The cut JCVI-syn3A YCp was then closed using the 102 base 380_40bp_161L, which has 40 base overlaps to both JCVISYN3A_0379 and JCVISYN3A_0381, and has a 22 bp *Mycoplasma gallisepticum* 161 CRISPR/Cas9 target sequence with a PAM (5'-GTATAAATACATCCAGGAGtgg-3' that had no homology elsewhere in JCVI-syn3A. This reaction was done in concert with the CRISPR/Cas9 genome cleavage as described above in Protocol I. The *M. gallisepticum* sequence in 380_40bp_161L oligonucleotide that joined the CRISPR/Cas9 cut JCVI-syn3A YCP while removing all of gene JCVISYN3A_0380 and parts of JCVISYN3A_0379 and JCVISYN3A_ 0381 put a new PAM in the genome that will be employed in the second round of CRISPR/Cas9. For the yeast transformation we mixed 100 ng of trp plasmid pCC1Bac-LC-YACTRP (GenBank accession number MN982904), 500-600 ng of guide RNA, 1 µg of 380_40bp_161L donor DNA and 100 µl of transformation competent yeast cells harboring the JCVI-syn3A YCp (VL648NCAS9_JCVI_Syn3A). After 2 hours of culture in YPDS media at 37°C, we plated 100 µl or 20 µl of culture on CM agar plates lacking histidine and tryptophan (CM Glucose Agar, Dry, w/o Histidine, Tryptophan; Teknova) with 2% agar. Colony PCR using was performed on 20 colonies using primers 380gRNA5, and 380_40bp_161L donor DNA. One fourth of colonies generated 303 bp amplicons indicating they lacked parts of genes JCVISYN3A_0379 and JCVISYN3A_ 0381 and all of gene JCVISYN3A_0380 (Supplementary Figure SA1A). We used our set of multiplex primers to assay JCVI-syn3A YCp/genome integrity to determine if any of the five positive clones appeared to be missing genome segments. All five clones appeared to have complete genomes (Figure SA1B). Clone #7, hereafter called clone 380_g5#7 (Figure SA1A) was chosen to use in subsequent steps.

The second round of CRISPR/CAS9 needed to create the needed point mutation cut the JCVI-syn3A YCp at the new *M. gallisepticum* PAM. The cut YCp was then re-circularized using a 1727 bp donor DNA. This donor DNA molecule contained the His230Ala mutation in gene JCVISYN3A_0380 expected to eliminate the gene 0380 protein's hydrolase activity. The second step CRISPR/CAS9 was performed on the first step positive clone 380_g5#7.

For the first step in the fusion PCR to build the 1727 donor DNA, we made a 558 bp PCR amplicon with the His230Ala mutation. The first step in this process was a PCR that used primers 50bp_380_F1, 380_R1 and JCVI-syn3A genomic DNA as template.  The 25 µl PCR was comprised of 12.5 μl 2X Qiagen multiplex master mix (Qiagen), 1 μl of each primer (0.2 μM final), 10.3 μl $H_2O$, and 0.2 μl (200 ng) JCVI-syn3A template. The PCR began with 2 minutes at 98 °C and then had 30 cycles of 98 °C for 10 seconds, 55 °C for 10 seconds, and 72 °C for 10 seconds, followed by 72 °C for 5 minutes. A 438 bp DNA band containing the His230Ala mutation was obtained for use in subsequent fusion PCR steps. The His230Ala mutation was introduced by primer 50bp_380_F1. The second step PCR used primers p380_F2 and 380_F(558) and JCVI-syn3A genomic DNA as template. The 25 μl PCR was comprised of 12.5 μl Q5® High-Fidelity 2X Master Mix (New England Biolabs), 1 μl of each primer (0.2 μM final), 10.3 μl $H_2O$, and 0.2 μl (200 ng) JCVI-syn3A template. The PCR began with 2 minutes at 98 °C and then had 30 cycles of 98 °C for 10 seconds, 55 °C for 10 seconds, and 72 °C for 10 seconds, followed by 72 °C for 5 minutes. A 188 bp DNA band was obtained as a donor DNA. Here, the His230Ala mutation was introduced via primer p380_F2. The above two PCR amplicons were column purified (Zymo Research). Using the two amplicons as template and 380_R1 and 380_F(558) primers and Q5 master mix (same PCR recipe and conditions as above) the 558 bp donor DNA with point mutation His230Ala in JCVISYN3A_0380 was obtained. Primers are listed in Supplementary Table SA8.

The next steps in the production of the mutagenic 1727 donor DNA molecule were done using a three-fragment fusion PCR. First, using primers 380_F4 and 380_R4, JCVI-syn3A genomic DNA as template a 402 bp DNA amplicon was synthesized comprising parts of genes JCVISYN3A_0380 and JCVISYN3A_ 0381. The 25 μl PCR was comprised of 12.5 μl PrimeSTARMax 2X Master Mix, 1 μl of each primer (0.2 μM final), 10.3 μl $H_2O$, and 0.2 μl (200 ng) JCVI-syn3A template. The PCR began with 2 minutes at 98 °C and then had 30 cycles of 98 °C for 10 seconds, 55 °C for 10 seconds, and 72 °C for 10 seconds, followed by 72 °C for 5 minutes. Second, using primers 380_F5 and 380_R2 and the same PCR recipe and conditions as just listed, a second 817 bp DNA amplicon was obtained. The above two PCR products were column purified (Zymo Research). The above 402 bp and 817 bp DNA fragments and the 558 bp donor DNA with the His230 Ala mutation were mixed with 380_R4 and 380_F5 primers. A PCR was performed using the aforementioned recipe and conditions to produce the 1727 donor DNA with point mutation His230Ala in JCVISYN3A_0380.

To produce a JCVI-syn3A YCp containing the His230Ala mutation in gene MMSYN_0380, we still needed to make another sgRNA that would function in a CRISPR/Cas9 reaction to cut the 380_g#7 yeast clone obtained previously at the *M. gallisepticum* PAM. To produce that sgRNA template, we did a PCR using *M. galli*_161_gRNA1, and gRNA_R as primers and the 83 base sgRNA_Template Ultramer as template as was done previously (Supplementary Table S8). The sgRNA was *in vitro* transcribed with the T7 RiboMAX™ Express Large Scale RNA Production System as done previously.

For the yeast transformation to produce a JCVI-syn3A YCp containing the His230Ala mutation we mixed 100 ng of URA plasmid pRS316 (ATCC® 77145™), 500-600 ng of guide RNA, 1 µg of 1727 bp mutagenic donor DNA, and 100 µl of transformation competent 380_g#7 yeast clone. After 2 hours of culture in YPDS media at 30 °C, we plated 100 µl or 20 µl of culture on CM agar plates lacking histidine and uracil. Primers 380_2jF(287) and 380_2jR(287) were used to screen positive colonies for the desired point mutation. 1934 bp DNA band was expected for positive colonies. A 310 bp DNA band was expected for wild type 380_g5#7. We screened 20 colonies and found five had the mutagenic 1727 donor DNA based on colony PCR (Figure SA1C). Multiplex PCR showed that all five positive colonies had complete JCVI_Syn3A YCps (Figure SA1D). Two positive colonies were Sanger sequenced for the 1727 bp region. Both colonies had the His230Ala mutation as designed. One of those colonies had an unwanted mutation. The correct yeast clone was used in a genome transplantation reaction as described previously to produce JCVI-syn3A bacteria with the His230Ala mutation in gene JCVISYN3A_0380.To make the control strain we generated a His230His (CAC to CAT) mutation in MMSYN1_0380 using the similar two-step process for CRISPR/CAS9 editing.

A c-t_1727 donor DNA was made by fusion PCR. Using primers c-T_380_F1 and 380_R4 and PrimeSTARMax master mix the first 759 bp DNA fragment was obtained. Using primers c-T_380_R1 and 380_F5 and PrimeSTARMax master mix the second 988 bp DNA fragment was obtained. His230His (CAC to CAT) in MMSYN1_0380 mutation was introduced via primers c-T_380_F1 and c-T_380_R1. The above two PCR product was column purified (Zymo Research). Using the above two DNA fragments as template and 380_R4 and 380_F5 primers and

PrimeSTARMax master mix the c-t_1727 donor DNA with point mutation His230His (CAC to CAT) in MMSYN1_0380 was obtained. The c-t_1727 donor DNA has a 40-bp overlap with the altered genome obtained from the first step.

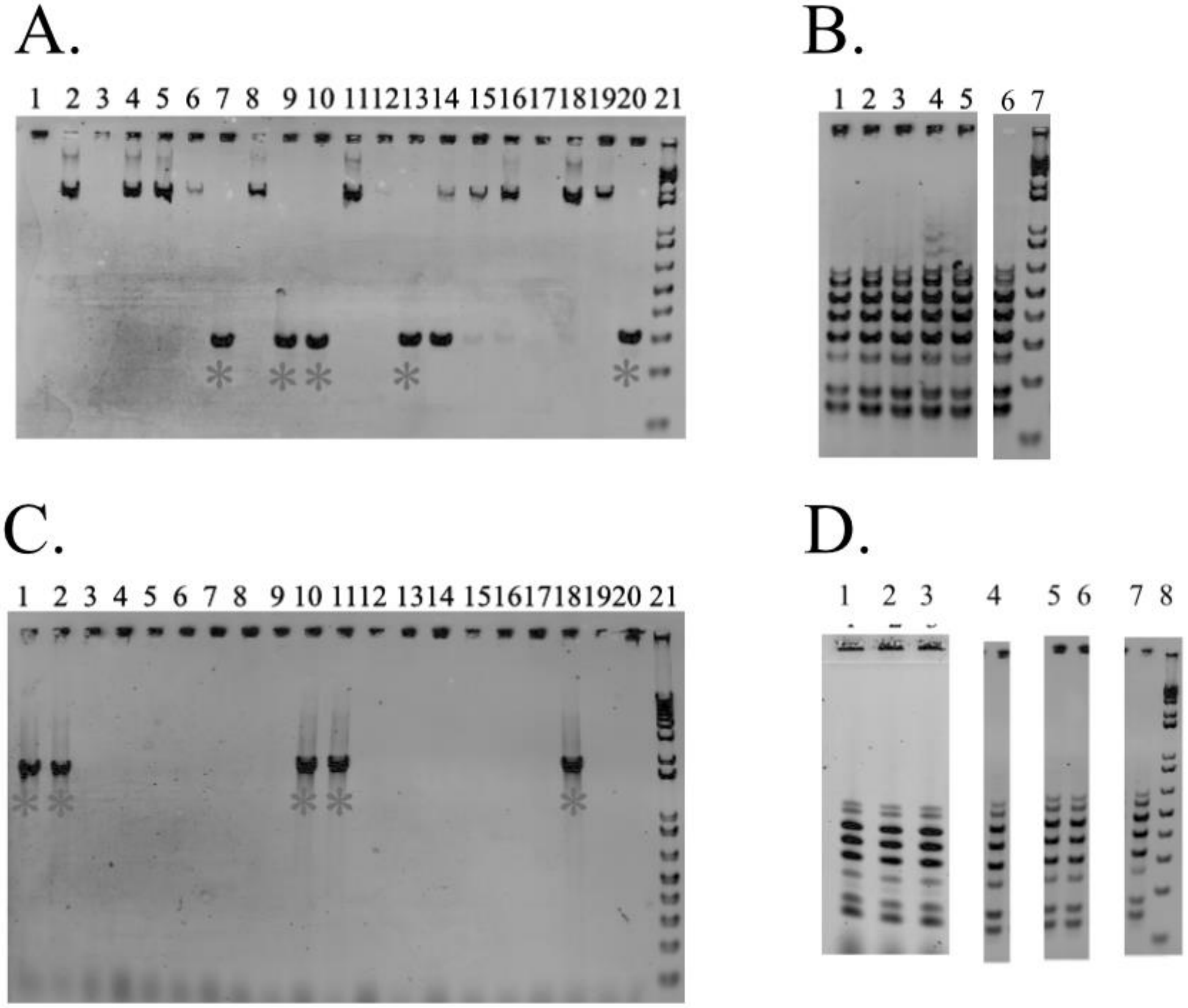

**Figure SA1. Colony PCRs for two step CRISPR/CAS9 with 1727 bp donor DNA to produce a JCVI-syn3A YCp with a His230Ala mutation in gene JCVISYN3A_0380.** (A) Colony PCR for 1st step CRISPR/CAS9 with 380gRNA5, and 380_40bp_161L donor DNA. Lane 1-20: colony PCR clone #1-20; Lane 21: 1kb Plus Ladder (ThermoFisher). Clones #7, 9, 10, 13, 20 showed the expected 303 bp bands for gene deletion.

(B) Multiplex primer PCR for clones in which gene 380 was deleted in (A). Lane 1-5: clones #7, 9, 10, 13, 20; lane 6: JCVI_Syn3A control; lane 6: 1kb Plus Ladder. All clones checked have complete JCVI_Syn3A genome. (C) Colony PCR for 2nd step CRISPR/CAS9 with M. galli_161gRNA1 and the 1727 mutagenic donor DNA as primers. Lane 1-20: colony PCR for clone 1-20. Clone # 1, 2, 10, 11, 18 have the expected 1934 bp DNA band for 1727 bp donor DNA insertion; lane 21: 1kb Plus Ladder. (D) Multiplex primer PCR for positive clones with ala_1727 donor DNA insertion. lane 1-3 and 5-6: multiplex primer PCR for clone # 10, 11, 18, 1, 2 with ala_1727 donor DNA insertion; Lane 4&7: **multiplex primer PCR for clone #19 and #7 with c-t_1727 donor DNA insertion; lane 8: 1kb Plus Ladder. * showed the positive colonies with gene deletion or insertion.**

**Supplementary Table SA1.** Primers for creating DNA molecules that serve as templates to produce synthetic gRNAs in a MEGAshortscript T7 transcription kit to be used for CRISPR/Cas9-based knockout of genes JCVISYN3A_0066, JCVISYN3A_0077, JCVISYN3A_0710, JCVISYN3A_0728, JCVISYN3A_0907, in the yeast centromeric plasmid containing a genome clone of JCVI-syn3A**.**

---

sgRNA_Template (83 bases)

GTTTTAGAGCTAGAAATAGCAAGTTAAAATAAGGCTAGTCCGTTAT
CAACTTGAAAAAGTGGCACCGAGTCGGTGCTTTTTTT

gRNA_R AAAAAAAGCACCGACTCGG

Forward primers (underlined regions are from genes to be deleted or modified)

0066_gRNA_F1

TAATACGACTCACTATAGGTATTAAAAGCTTGTACAACG
TTTTAGAGCTAGAAATAGCAA

0066_gRNA_F2

TAATACGACTCACTATAGGTTATCAATTTGTAATAAGCGT
TTTAGAGCTAGAAATAGCAA

0077_gRNA_F1
TAATACGACTCACTATAGGCAAATATTACAGCAACTAAG
TTTTAGAGCTAGAAATAGCAA

0077_gRNA_F2
TAATACGACTCACTATAGGTATTAAATAAATCTGCTCTGT
TTTAGAGCTAGAAATAGCAA

0710_gRNA_F1
TAATACGACTCACTATAGGTTTAACTGCAGATCAAGGGT
TTTAGAGCTAGAAATAGCAA

0710_gRNA_F2
TAATACGACTCACTATAGGTAGAAGCGATTAAATTTGCG
TTTTAGAGCTAGAAATAGCAA

0728_gRNA_F
TAATACGACTCACTATAGTATTAATATTAGATATGGATGT
TTTAGAGCTAGAAATAGCAA

0907_gRNA_F1
TAATACGACTCACTATAGGTAAAGGTCAAGCGATTAAGT
TTTAGAGCTAGAAATAGCAA

---

**Supplementary Table SA2. Single donor DNA patches for yeast homologous recombination-based repair and target gene removal or mutation of JCVI-syn3A YCPs after CRISPR/Cas9 mediated cleavage.**

---

100 bp patch for 0066
ACACCATCTTTATCATTTGAATAATCGGTGATATCATCAGCTATTTCTTTATTAATAT
TCTATAGCATTTTAACTTTATTTCAAGCATATATTAAAAAAA

100 bp patch for 0077
ATTTATATTTGTATCAGTGATATATTTTGCTTTTTGCTTAATTGGATCAATTACTACTC
CTTAATAATCTTTCTTTTTAAAATTATTATTTGTTATTTAA

100 bp patch for 0710

GAAATAAAACAATAATTCAAAATACTAAAAACTGAAAAAATACTAATTAAAAAAAC
TCTCTTTTCTATTATTATGATTATAACAAAATCAATTATTCTAA

100 bp patch for 0728

AGTGATTATTTAGGTTTAGTTAACTATAATGAATAAACCAGAAATTAAGTCAGTTGA
AGAATTTATTAAACACTATATACTAGGAGAATAAAAATGAAAG

100 bp patch for 0907

AATATTTTTATTATGTTCATGTTTACAAATATAAGAGTTATCATAAAACAACTAGCA
CCTACAATATTATCTTAATTTATTTATATTCTTTTAAATACAA

100 bp patch for 0380

CTTAAAAAAGAAAAACAAAATCAGAATATTTAAATAATTTAGGTGCAATATTTTTGT
GCAGATAAGATTAGTATTGAAAGAGATTATTTAGGTGTTGATA

**Supplementary Table SA3. Primers for yeast colony PCRs to determine if CRISPR/Cas9-based deletion of genes succeeded or failed (reverse primer or primers that would produce a band if the gene were still present are in italics). The number in parentheses on the right of each primer name is the size in base pairs of the PCR amplicon that would or wound not be produced.**

| | |
|---|---|
| 66 LJ F(239) | TTGTCAAATTGTTCTTTTGAAGTGA |
| 66 2J R(239) | GAGAAGATAGCAAGCTGCCAGT |
| *66 LJ R(323)* | *TGTTTGATATTTCAGTTAAAGGTGTTG* |
| 77 LJ F(499) | TCATAAATGAAACAAATTTACCTTGTC |
| 77 2J R(499) | GCAGCTAGTTTTCAATATACAGCA |
| *77 LJ R(620)* | *CAGCAACAGCTAGTGATGTTTATTG* |

710 LJ F(555) AAAATAAGGGGAAAATAAAAGGTT
710 2J R(555) TGATCTACGGGCGCATATAAA
*710 LJ R(286) GATCAAGGTGGAGTTGGAGAA*

Ljun_F(272) TGCCAAATATTGATGGAGCTT
Ljun_R(272) TGATCAACTAACAAACCGTGATG
*2Sjun_F(494) CCAATTTGAGCAATTGGTACTG*
*2Sjun_R(494) CCCATTTGATTATCAGGCAGA*

907 LJ F(147) TTTCTTAACGTATTATCAAAATCTGAA
907 LJ R(147) CGTCAAATGGAAAGAATGAATTT
*907 2J F(415) CCTAAATCAATTACAGGTTCAACTCC*

380 LJ F (364)TTGAATCATAGCTACAGTTTTTGGA
380 junc R (445) AAGCTGGAATTAATCAAACTTCA
380 2j R (604) AAACAACAACTAGCATTTACTAATCCA

**Supplementary Table SA4. Multiplex primer pairs for diagnostic PCR to assess JCVI-syn3A genome and YCp integrity and PCR amplicon sizes resulting from each primer pair.**

| Name | Sequence | Amplicon length |
|---|---|---|
| RCO1776 | ACCAAGTACAATGCTAAGTG | 150 bp |
| RCO1777 | CTCCTGAATACTTATTTGAAAAAC | |
| SBIR-2F | TTTTAGTATGTTTAAGGAGAAGA | 186 bp |
| MPCR9-2R | TGATGGTGCATAACGTAATC | |
| MPCR11-3F | TGATGATGATAATAAATTAAATCCT | 267 bp |
| RCO1778 | AAAAAGCTATTTTTTACAAGTTCAA | |
| RCO1779 | TATTAGGATCAGTAGCTAAAGG | 330 bp |
| RCO1780 | GTTTAAGTCAATCTTTTCTTCTTG | |
| RCO1781 | ACCTATTATTTATAAATAACAATGC | 409 bp |

| | | |
|---|---|---|
| RCO1782 | TTGTAGCAACTGGATATAATGG | |
| SBIR-6F | GTTTTACCAACTCCAGGTTC | 496 bp |
| LJ 6R(496) | GTGTTAAACAAGGATATTGTG | |
| RCO1805N | AACTCCTATGGGTGTGTTG | 583 bp |
| RCO1784 | TAAATTATTATGACAATTCACTTTC | |
| RCO1785 | GACTATTGCCTCCAATTAGA | 650 bp |
| RCO1786 | TCTTTTGCTAATTGTTGACTTG | |

**Supplementary Table SA5. Primers to construct sgRNA templates to cleave the JCVI-syn3A YCp downstream of the HIS3 yeast selectable marker prior to insertion of new copies of the target genes.**

sgRNA_Template (83 bases)
GTTTTAGAGCTAGAAATAGCAAGTTAAAATAAGGCTAGTCCGTTATCAACTTGAAAAAGTGGCACCGAGTCGGTGCTTTTTT
His_gRNA_primer F
TAATACGACTCACTATAGAAAAAAATATAGAGTGTACTGTTTTAGAGCTAGAAATAGCAA
gRNA_R
AAAAAAAGCACCGACTCGG

**Supplementary Table SA6. Primers for knocking in gene JCVISYN3A_0066, JCVISYN3A_0077, JCVISYN3A_0710, and JCVISYN3A_0907 in the landing pad using CRISPR/CAS9.**

**<u>First PCR round primers to be used with JCVI-syn3A genomic DNA template</u>**

lox66 F1
GTTGATATATTATAGTATAGACCTACCGTTCGTATAAGAAACCATATACGAAGTTATTATATTTCTCCTAGTCTAT

lox66 R1

GCTGTTAATTCTGATTCAGTACCGTTCGTATAGCATACATTATACGAAG TTATTAAAAAAGAACCTTAATGG

lox77 F

GTTGATATATTATAGTATAGACCTACCGTTCGTATAAGAAACCATATAC GAAGTTATCTAATTTAAAATATATTTTTCTATAAATTTACC

lox77 R

GCTGTTAATTCTGATTCAGTACCGTTCGTATAGCATACATTATACGAAG TTATTGAGTTATTGTACAGATAATG

lox710 F

GTTGATATATTATAGTATAGACCTACCGTTCGTATAAGAAACCATATAC GAAGTTATAAAATAAGGGGAAAATAAAAGGTT

lox710 R

GCTGTTAATTCTGATTCAGTACCGTTCGTATAGCATACATTATACGAAG TTATTGATCTACGGGCGCATATAAA

lox907 F

GTTGATATATTATAGTATAGACCTACCGTTCGTATAAGAAACCATATAC GAAGTTATTTAGTATTCAATCTCTCAA

lox907 R

GCTGTTAATTCTGATTCAGTACCGTTCGTATAGCATACATTATACGAAG TTATATTACAAATTATTCAAACTTATGC

**Second PCR round primers to be used with 1st PCR round amplicon templates**

his land F

GAAAAAAAAAAATGAAAATCATTACCGAGGCATAAAAAAATATAGAGTGTACTAGT TGATATATTATAGTATAGACC

ARS land R

TTCCATCATTAAAAGATACGAGGCGCGTGTAAGTTACAGGCAAGCGATCCGCTGTTA ATTCTGATTCAG.

**Supplementary Table SA7. Primers for yeast colony PCRs to determine if CRISPR/Cas9-based deletion of genes succeeded or failed (reverse primer or primers that would produce a band if the gene were still present are in italics). The number in parentheses on the right of each primer name is the size in base pairs of the PCR amplicon that would or wound not be produced**

| | |
|---|---|
| 2junc F(166) | TGTGATAATGCCAATCGCTAA |
| 734TC-F(358) | GTCATCCGCTAGGTGGAAAA |
| lox junc F(232) | TGCTATACGAACGGTACTGAATC |
| lox junc R(232) | TGTAAAGTACGCTTTTTGTTGAAATTAATGG |
| 66 LJ R(323) | TGTTTGATATTTCAGTTAAAGGTGTTG |
| 77 LJ R(620) | CAGCAACAGCTAGTGATGTTTATTG |
| 710 LJ R(286) | GATCAAGGTGGAGTTGGAGAA |
| 907 LJ R(478) | TCAAAATACTTGAGATCAAATTGAAAA |
| Ljun_R(272) | TGATCAACTAACAAACCGTGATG |

**Supplementary Table SA8. Primers used to mutate 0380 codon 230 from His to Ala. Lower case bases are changes from the JCVI-syn3A gene 0380 wild type sequence.**

Guide RNA template synthesis primers

161_gRNA_F1

TAATACGACTCACTATAGGTATAAATACATCCAGGAGGTTTTAGAGCTAGAAATAGCAA

380gRNA_F5

TAATACGACTCACTATAGgCTGTAGCTATGATTCAAAAGTTTTAGAGCTAGAAATAGCAA

*M.galli*_161_gRNA1

TAATACGACTCACTATAGGTATAAATACATCCAGGAGtGGTTTTAGAGCTAGAAATAGCAA

sgRNA_Template (83 bases)

GTTTTAGAGCTAGAAATAGCAAGTTAAAATAAGGCTAGTCCGTTATCAACTTGAAAAAGTGGCACCGAGTCGGTGCTTTTTT

Donor DNA molecules and 1727 bp mutagenic donor DNA synthesis primers

380_40bp_161L

AAAAAATTTTAAACCAATTGAAGTTTTTGGAATAGCTATTGTATAAATACATCCAGGAGtggTTTAGTTATTCTTATGGTCAAATTCCAAAATTACCTAAAT

50bp_380_F1

GCTCATGATTTAGCTATTAAATGAAATGTTGATCCTAAAAAAGCCTTAATAGCTGGAACACTTGCAGA

| | |
|---|---|
| 380_R1 | CATTGATCCAATATTATCTTTTCCA |
| 380_F(558) | GACAAACAAATCAAAGAAATTAATGA |
| 380_R4 | ATTTAGGTAATTTTGGAATTTGAC |
| 380_F4 | TGGAAAAGATAATATTGGATC |
| 380_F5 | AAAAAATTTTAAACCAATTGAAGTTTTTGG |
| 380_R2 | CATTAATTTCTTTGATTTGTTTGTC |
| c-T_380_F1 | AACACTTCAtGATATTACAAAAAG |
| c-T_380_R1 | TTGTAATATCaTGAAGTGTTCCAGCTATTAAG |
| 380_R4 | ATTTAGGTAATTTTGGAATTTGAC |
| 380_F5 | AAAAAATTTTAAACCAATTGAAGTTTTTGG |

Yeast colony PCR molecules to identify His230Ala mutants in MMSY1_0380

| | |
|---|---|
| 380_2jF(287) | AAAGCAAAATAATGATCCTGAACT |
| 380_2jR(287) | AATAAAAACATCTCCACTTGCAATA |
