## Supplementary material for "Metabolite Damage and Damage-Control in a Minimal Genome": Sup Data 5: S5F-Instructions for recreating Figure 7.docx

**Introduction**

The escher map in Figure 7, while useful as a broad overview, lacks the detail for full analysis. However, once rendered in the escher web-tools, this map can be zoomed, expanded, modified, and explored in much greater detail. Unfortunately, it is a multi-step process to load the map into the escher interface. This document will walk you through that process.

**Instructions**

1. Go to the escher website run by the Palsson lab: <https://escher.github.io/#/>
2. Don’t change any settings on the homepage – just click on the “Load map” button
3. Your screen should now look like this:

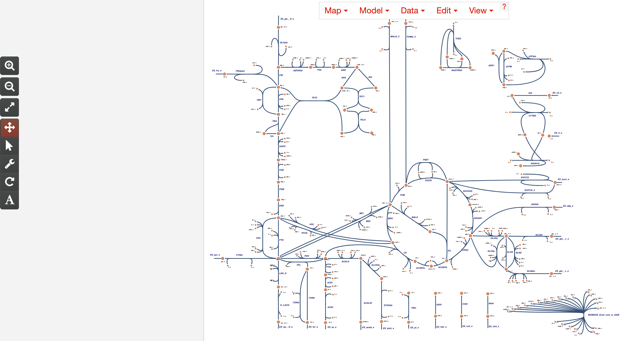

4. Click on the “Map” dropdown on the far left of the control panel at the top of the screen, and select the “Load map” option.

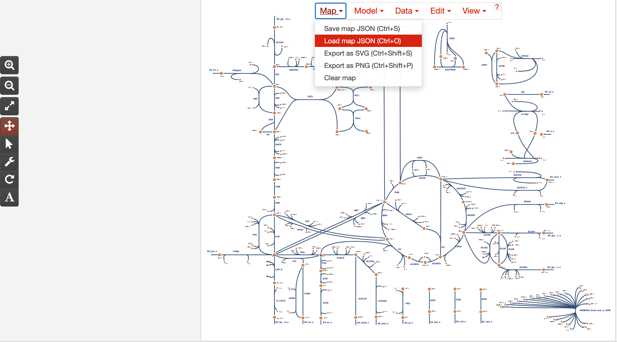

5. Select the supplementary file “S5C-JCVISyn3Exp_map.json” as your map to load and submit. Your screen should now look like this:

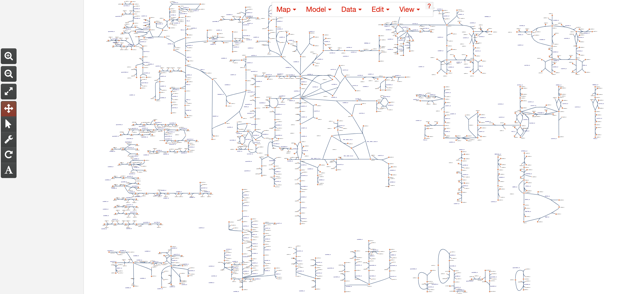

6. Now, click on the “Model” dropdown on the control panel at the top of the screen, and select the “Load COBRA model JSON” option. Select the supplementary file “S5B-JCVISyn3Exp_model.json” as your model to load and submit. The screen should not change appreciably once you do this.
7. Return to the “Model” menu, and now click on the “Update names and gene reaction rules using model” button. Again, the screen will not change much, but a message will show up at the bottom saying “Successfully converted attributes”.
8. Now, click on the “Data” dropdown in the panel at the top of the screen, and select the “Load reaction data” option. Select the supplementary file “S5D-Flux data.csv” as you data to load and submit. Now reactions should change color, but the color scheme will be different from Figure 7:

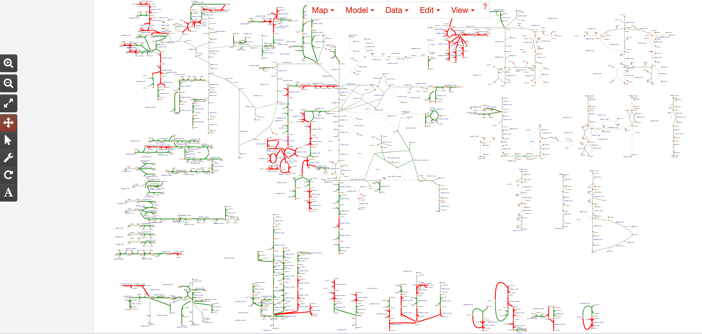

9. Note, the fluxes you just loaded are not the “true” fluxes from the FBA study. Those fluxes are provided in “Supplemental_data_S4_Modeling_data.xlsx” in the S4B tab. The values in the flux file have been adjusted to permit coloration of reactions depending on whether they were in the original model (flux ~10), in the ModelSEED database (flux ~1000), predicted spontaneous reactions (flux ~100), and predicted promiscuous enzymatic reactions (flux ~10000). To replicate the color scheme in Figure 7, click on the “View” dropdown in the panel at the top of the screen, and select the “Settings” option. In the “View and build options” region, click on the “Descriptive names” radio button next to the label “Identifiers”. In the “Reactions” region of the control panel that comes up, click on the gray bar on the far left of the color bar, and change the number to “10” and the color to magenta. Click on the gray bar on the far right, and change the number to 10000 and the color to blue. Click anywhere on the color bar between the two gray bars, and a new gray bar will appear. Click on this gray bar and change its number to 100 and its color to red. Finally, click on the color bar again between all the existing gray bars, and another new gray bar will appear. Change the number in this bar to 1000 and the color to green. The screen should now look like this:

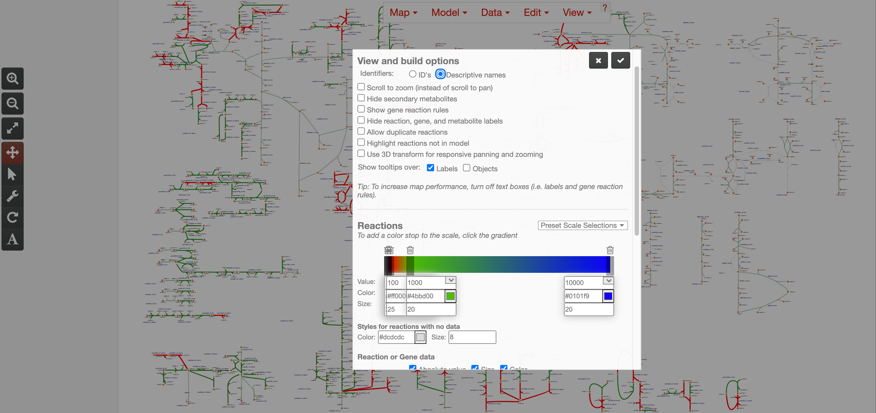

10. Click the check box at the top of the “Settings” control panel to accept the changes. The screen should now look like this:

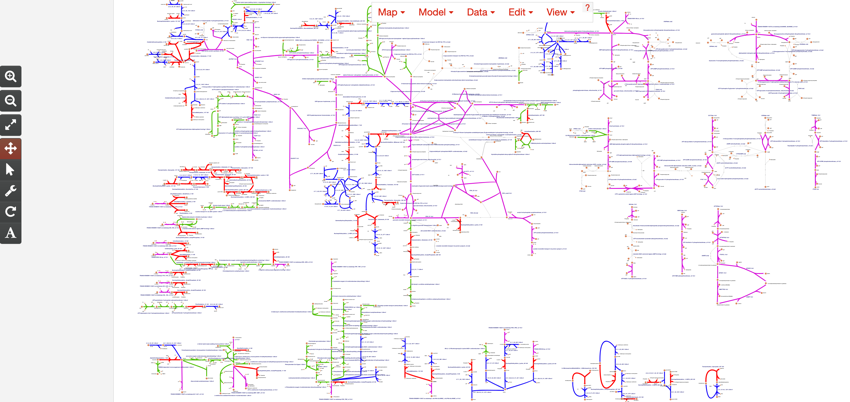

11. Finally, click on the “Data” dropdown in the panel at the top of the screen again, but this time, select “Load metabolite data” and load supplementary file “S5E-Metabolite data.csv”. This will paint metabolites with metabolomics data. Your screen should now look like this:

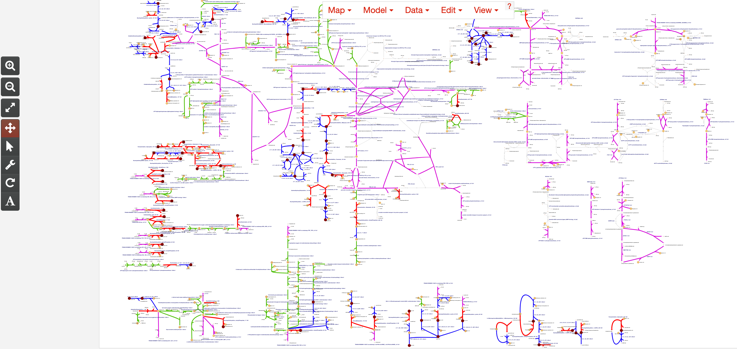

And with that, we have successfully replicated Figure 7 in a fully functioning escher map, which you can then explore on your own in whatever level of detail you prefer.
