## Supplementary material for "Metabolite Damage and Damage-Control in a Minimal Genome": SupplemtalTablesFigsMethods

### Supplemental Methods and Tables

### Supplemental Methods

#### Compilation of functional annotations for Mycoplasma JCVI-Syn3A proteins

The Mycoplasma JCVI_Syn3 genome was imported in the SEED database (1) [https://pubseed.theseed.org/?page=BrowseGenome&organism=1806462.19] to capture the SEED annotations , these were exported and compared with the curated sets of annotations of both Mycoplasma JCVI_Syn3 and Mycoplasma JCVI_Syn3A generated by the Danchin (2), the Stülke (3) and the Zhang (4) groups respectively to produce a manually curated composite annotation, that were colored in different functional categories to allow easy sorting (Supplemental Data S1) . The essentiality data was added from our previous studies (5).

#### Protein expression and purification and enzyme assays

- **CoA disulfide reductase**

The pET28a derivative encoding CoADR (pET28-887) was transformed into BL21 CodonPlus (DE3)-RIPL (Stratagene) cells). Cultures (250 mL) were grown at 37 °C, 300 rpm, in LB medium containing 50 µg/mL kanamycin, 34 µg/mL chloramphenicol. When an OD600 of 0.6 was reached, the incubation temperature was reduced to 21 °C and IPTG was added to a final concentration of 1 mM, as well as ETOH to 4% and cultures were incubated for another 16 h. Cells were then harvested by centrifugation (6,000 *g*, 10 min, 4°C), resuspended in 50 mM sodium phosphate, 300 mM sodium chloride, 10 mM imidazole, 1 mM FAD, pH 8.0 and sonicated. Debris was removed by centrifugation at (14,600 *g*, 10 min, 4°C) and 1mL of HisPur™ Ni-NTA Superflow Agarose (ThermoFisher Scientific) slurry was added and the solution was incubated for 30 min at 4 °C. The suspension was then transferred to an empty polypropylene column and washed with 50 mL of 50 mM sodium phosphate, 300 mM sodium chloride, 1mM FAD, 20 mM imidazole, pH 8. Purified CoADR was then eluted with 2 mL of 50 mM sodium phosphate, 300 mM sodium chloride, 1mM FAD, 250 mM imidazole, pH 8.0. The eluate was then desalted with a PD-10 column (GE Healthcare) equilibrated with 50 mM potassium phosphate pH 8.0, 10% glycerol, pH 8, flash frozen in 100 µL aliquots and stored at -80 °C until needed.CoADR activity was measured spectrophotometrically by monitoring

NAD(P)H consumption by absorbance at 340 nm ($A_{340}$) every 10-15 s on a Beckman DU 7400 spectrophotometer. These 100 µL assays contained 50 mM potassium phosphate pH 8.0, 0.2 mM NAD(P)H, 1 mM EDTA-KOH pH 8.0, 0.5 mM co-substrate and 2.86 µg of CoADR. As CoADR oxidizes NAD(P)H without the addition of a co-substrate, this basal rate was subtracted from other measurements. To determine the pH optimum, the pH of the potassium phosphate buffer was varied from 6.4-8.2. To determine the kinetic constants with respect to CoAD consumption, the concentration was varied from 3-1500 µM.

- **NadD-YqeK**

The pET28a derivatives encoding JCVISYN3A_0380, $NadD_{Bs}$, JCVISYN3A_0380H233A, and the YqeKYqeK domain of JCVISYN3A_0380) ($YqeK_M$) were electroporated into BL21-CodonPlus (DE3)-RIPL cells (Stratagene). Cultures (0.5 liter) were grown at 37°C in LB medium containing 50 µg/ml kanamycin, 50 µg/ml spectinomycin, and 34 µg/ml chloramphenicol. When A600 reached 0.6–0.8, isopropyl β-D5 1-thiogalactopyranoside was added to a final concentration of 1 mM and incubation was continued for 4 h at 22°C. Cells were harvested by centrifugation (6300 g, 15 min), resuspended in 50 mM Tris-HCl, 300 mM NaCl, and 10 mM imidazole, pH 8.0, and sonicated. The lysate was centrifuged at 7000 g for 15 min and the supernatant was incubated with 1 ml of $Ni^{2+}$-NTA 50% slurry (Qiagen) for 20 min at 4°C. The slurry was poured into a column and allowed to drain by gravity. After washing with 50 ml of 50 mM Tris-HCl, 300 mM NaCl, and 20 mM imidazole, pH 8.0, proteins were eluted with 2.5 ml of this buffer containing 250 mM imidazole, desalted on PD-10 columns (GE Healthcare) pre-equilibrated in 20 mM HEPES, pH 7.2, 100 mM NaCl, 0.2 mM DTT, 10% (v/v) glycerol. Proteins was desalted into 20 mM HEPES, pH 7.2, 100 mM NaCl, 0.2 mM DTT, 10% (v/v) glycerol using a PD-10 column, aliquoted, flash-frozen in liquid N2, and stored at -80°C. Protein purity was confirmed by SDS-PAGE.

NadD assays contained 0.5 mM NaMN, 4 mM $MgCl_2$, 5 u/ml yeast inorganic pyrophosphatase, 1 mg/ml BSA and 2 mM NTP (dATP, CTP, GTP or UTP) in 20 mM HEPES, pH 7.2, 100 mM NaCl, 0.2 mM DTT, 1% glycerol in 100 µl final volume. (YqeK assays) contained 0.5 mM $NaAD^+$ or $NaAD^+$ analog, 2 mM $MgCl_2$, 1 mg/ml BSA and 0.2 u CIP in the same buffer and a total volume of 20 µl. Use of CIP was necessary to resolve product peaks chromatographically. Assays were started with addition of protein, NadD assays were stopped with 1 volume acetone. Assays were dried *in vacuo*, redissolved in 100 µl water and passed through Amicon Ultra 0.5 ml 10K units. YqeK assays were stopped by adding 100 μl water to the reaction mixture and passed through Amicon Ultra 0.5 ml 10K units. A 5 μl aliquot of the NadD assay flow-through, or a 40 μl aliquot of the YqeK assay flow-through was analyzed by HPLC with UV detection (Waters 2695 Separation module and Waters 2998 PDA detector) using a C18 column (Thermo Scientific Hypersil GOLD C18, 5 μm, 250×4.6 mm) with a column guard. The following gradient of buffer A (10 mM $KPO_4$, pH 6.0) and B (acetonitrile) was used: 0-10 min, gradient from 100% A, 0% B to 99.5% A, 0.5% B; 10-20 min gradient to 90% A, 10% B; 20-22 min isocratic at 90% A, 10% B; 22-23 min gradient to 100% A, 0% B; 23-38min isocratic at 100% A, 0% B.

- **HAD proteins**

Plasmid pET28b carrying HAD genes (JCVISYN3A_0066, JCVISYN3A_0077, JCVISYN3A_0907, or JCVISYN3A_0728) were transformed to *E. coli* BL21-CodonPlus (DE3)-RIPL (Stratagene) cells by electroporation. Cultures (0.25 l) were grown at 37 °C with vigorous shaking in LB medium containing 50 μg/ml kanamycin, 34 μg/ml chloramphenicol and 50 μg/ml spectinomycin until $A_{600}$ reached around 0.6. IPTG was added (final concentration 1 mM) to each culture and incubation was continued for 4 h at 37 °C. Cells were harvested by centrifugation (4,000*g*, 20 min), resuspended in lysis buffer (50 mM $NaH_2PO_4$, 300 mM NaCl, and 10 mM imidazole, pH 8.0), and sonicated. The extract was clarified by centrifugation at 10,000g for 30 min at 4°C. The supernatant was incubated at 4°C with a 1 ml $Ni^{2+}$-NTA 50% slurry (Qiagen) for 1 h and poured into a column, allow to drain. The column was washed with 16 ml washing buffer (50 mM $NaH_2PO_4$, 300 mM NaCl, and 20 mM imidazole, pH 8.0). Proteins were eluted with 2.5 mL elution buffer (50 mM $NaH_2PO_4$, 300 mM NaCl, and 250 mM imidazole, pH 8.0) and desalted on PD-10 columns (GE Healthcare) equilibrated in 50 mM Tris-HCl, pH 7.0, 5 mM $MgCl_2$, 10% (v/v) glycerol.

Purified HAD proteins were tested for general phosphatase activity against *p*-nitrophenyl phosphate (pNPP) as substrate using BIOMOL green (Enzo Life Science, Farmingdale, NY, USA) according to manufactures instruction (the assays were run for 15 min at 37°C and contained 1 mM PNPP and 1μg of enzyme). The buffer, pH and metal were optimized for each HAD protein against pNPP. Purified HAD proteins were then screened for natural substrate against a panel of 94 phosphorylated metabolites (Table S1) using Malachite Green reagent as previously described (6). The screening assays contain 50mM pH 5.9 MES-KOH, 2mM $MgCl_2$,

0.67 μg enzyme and 0.25 mM substrate for JCVISYN3A_0728, 50mM pH 5.5 MES-KOH, 5 mM $MgCl_2$, 1.86 μg enzyme and 0.25 mM substrate for JCVISYN3A_0907 and 50 mM pH 5.5 MES-KOH, 5mM $MgCl_2$, 3 μg enzyme and 0.25 mM substrate for 0066, and JCVISYN3A_0077).

- **JCVISYN3A_0400 (DJ-1) protein**

The pET15b plasmid expressing JCVISYN3A_0400 was transformed into BL21(DE3) *Escherichia coli* (Novagen). Cells were grown in Luria-Bertani (LB) medium supplemented with 100 μg/mL ampicillin at 37 °C with shaking until the $OD_{600}$ reached 0.5–0.7. Protein expression was induced by the addition of isopropyl β-d-1-thiogalactopyranoside (IPTG) to a final concentration of 0.5 mM and the culture was grown with shaking for four hours. Cells were harvested by centrifugation and the recombinant JCVISYN3A_0400 protein was purified as previously descried (7). Purified JCVISYN3A_0400 in storage buffer (25 mM HEPES pH=7.5, 100 mM KCl, 2 mM DTT) gave a single band on overloaded Coomassie-stained SDS-PAGE gels. Protein concentration was determined by $A_{280}$ using a calculated extinction coefficient of 15,930 $M^{-1}$ $cm^{-1}$ (ExPASY), snap-frozen in 50-100 μL aliquots in liquid $N_2$, and stored at -80°C until needed.

The methylglyoxalase activity of JCVISYN3A_0400 was measured using a coupled assay of L-lactate oxidase and Amplex Red/horseradish peroxidase (HRP). In this assay, the methylglyoxalase activity DJ-1 generates exclusively L-lactate (8). which is oxidized by L-Lactate oxidase and molecular oxygen to generate pyruvate and $H_2O_2$. The liberated $H_2O_2$ and Amplex Red are co-substrates for HRP, generating fluorescent resorufin. The rate of $H_2O_2$ generation was monitored using the Amplex Red Hydrogen Peroxide/Peroxidase Assay Kit (Invitrogen). Purified methylglyoxal (MG) was the gift of Dr. John Termini (City of Hope, Los Angeles, CA, USA). Various concentrations of MG were added to the Amplex Red working solution (including Amplex Red reagent and horseradish peroxidase (HRP)), 0.4 U L-lactate oxidase (Sigma L9795), in reaction buffer (50 mM sodium phosphate, pH 7.4). The reaction was then initiated by adding final concentration of 0.1μM JCVISYN3A_0400. For control experiments, human DJ-1 (0.1μM) and *Saccharomyces cerevisae* Hsp31 (0.05 μM). $H_2O_2$ generation was measured for five minutes using a Cary Eclipse spectrofluorimeter (Varian, Palo Alto, CA, USA) at 22°C with an excitation wavelength at 540nm and emission wavelength of 590nm. All reaction rates were linear over this timeframe, and the slope of the best-fit line was converted to $H_2O_2$ concentration using a standard curve generated with known concentrations of $H_2O_2$. The generation of $H_2O_2$ was linear in JCVISYN3A_0400 concentration and did not change appreciably when the L-lactate oxidase concentration was increased, establishing that the measured rates reflect the methylglyxoalase activity of JCVISYN3A_0400.

For deglycase activity measurement, JCVISYN3A_0400 protein was dialyzed against degassed PBS buffer pH 7.4 (10 mM $Na_2HPO_4$, 1.8 mM $KH_2PO_4$, 137 mM NaCl, 2.7 mM KCl) immediately prior to the assay to remove DTT which might interfere with generation of the hemithioacetal substrate. 10 mM coenzyme A (CoA; MP Biomedicals) and 10 mM of MG (gift of Dr. John Termini, City of Hope, Los Angeles, CA, USA) were mixed and incubated at 25°C until the $A_{288}$ value was stable (typically two hours), indicating that MG-CoA hemithioacetal formation was complete. Concentrations of MG-CoA hemithioacetal ranging from 0.1 mM to 4 mM were incubated in the cuvette for additional 10 minutes before JCVISYN3A_0400 was added to a final concentration of 10 μM. $A_{288}$ values were then measured for six minutes and the slope of the linear decrease was determined. All measurements were repeated at least three times. A buffer-only baseline was measured in order to correct for spontaneous (*i.e.* uncatalyzed) loss of 288 nm signal, which is minor. The difference between the buffer-only and buffer-enzyme curve was taken to reflect enzyme activity. The absorption change was converted to the change in MG-CoA hemithioacetal molar concentration by using the extinction coefficient of the related MG-N-acetyl-cysteine hemithioacetal of 98 $M^{-1}cm^{-1}$ (9).

**Metabolomics analysis of JCVI-Syn3A derivatives**

All JCVI_Syn 3A isolates were grown in SP4-KO medium and were harvested during of log-phase growth. Cell yields were determined prior and at the time of harvest as described in detail in Supplemental_data S2. Three biological replicates of each strain and three technical replicates from each biological sample were distributed and further pelleted/rinsed/flash frozen/stored. Cell pellet samples were fully randomized through sample extraction, analytical analysis, and data processing. 250 µL of 3:3:2 (isopropanol/acetonitrile/water) (v/v/v) was added to the original tube prepared for metabolomics analysis, vortexed vigorously for 30 seconds, sonicated for 1 minute, and all contents was transferred to a new clean 2mL round-bottomed microcentrifuge tube. An additional 250 µL of 3:3:2 was added to the original tube, vortexed, sonicated, and also transferred to the round-bottomed 2mL microcentrifuge tube to maximize recovery. Two three mm stainless steel balls were added to the round bottom microcentrifuge tube and processed in Geno/Grinder tissue homogenizer (Spex SamplePrep; Metuchen, NJ) at 1500 rpm for 1 minute. The tube was then centrifuged at 14000 rcf for 3 minutes, 450 µL of supernatant was transferred to a clean 1.5 mL microcentrifuge tube that was dried under vacuum overnight. Dried samples were stored at -80°C until analysis. Eight method blanks were generated by starting with 50 µL of methanol and no cell pellet, and these blanks were extracted evenly distributed throughout experimental sample extractions. A pooled quality control (QC) sample was created by transferring 50 µL of the remaining supernatant after centrifugation and removal of 450 µL of extract from each sample. This pooled QC was vortexed thoroughly to mix, and 450 µL of QC mix was transferred into eight 1.5mL microcentrifuge tubes, dried and stored in the same manner as samples. The LC-MS grade water, acetonitrile, and methanol used in metabolomics analysis were purchased from Fisher Scientific (Waltham, MA), and isopropanol, formic acid, and ammonium formate were purchased from Sigma-Aldrich (St. Louis, MO).

Prior to analytical analysis, dried samples were resuspended in 100 µL 8:2 (acetonitrile/ water) (v/v) with internal standards including 5 µg/mL Val-Tyr-Val and 42 other measured exogenous compounds (Supp Table 3B), vortexed for 30 seconds, sonicated for 1 minute, centrifuged at 14000 rcf for 2 minutes, and transferred to amber screw top vials with micro-insert for LC-MS/MS analysis. Analytical analysis was performed using a Vanquish Focused UHPLC coupled to a Q-Exactive HF mass spectrometer (ThermoFisher Scientific) as in Niehaus et al. (10). In summary the analysis used a Waters Acquity UPLC BEH Amide column (150mm x 2.1 mm id, 1.7 µm particle size) coupled to Acquity UPLC BEH Amide VanGuard precolumn (5 x 2.1 mm; 1.7µm) (Waters, Milford, MA) with mobile phases of LC-MS grade water, and 95:5 (v/v) acetonitrile/water, each at 10mM ammonium formate and 0.125% formic acid. Mass spectra were collected in data dependent mode with top four ions from each MS1 scan being selected for MS/MS fragmentation. Electrospray ionization analysis was conducted using a spray voltage of positive 3kV and negative 3kV as separate analyses. Three microliters of sample were injected for positive mode ionization, and 5 µL injected for negative mode analysis. Blanks and QC samples were analyzed evenly spaced throughout all analyses. Additionally, a pooled quality control sample was measured with a mass scan range window of 100 m/z sequentially 9 times to cover m/z 60-900 in order to increase data dependent MS/MS spectra coverage for use in compound identification.

Data was processed using open source software MS-DIAL (11) version 3.70. MS-DIAL performed baseline correction, deconvolution, peak detection, alignment, gap filling, adduct identification, m/z-RT library matching, and MS/MS library matching. MS-DIAL parameters are reported in Table S2. An in-house m/z-RT library developed from authentic standards was matched to peaks identified by MS-DIAL, and MS/MS spectra were matched to library spectra from the Mass Bank of North America (MoNA), and NIST17. Manual inspection of each annotated compound was conducted to confirm m/z-RT library match, and/or MS/MS library match which are reported in supplementary data S3. Peak height was reported as intensity for each annotated metabolite in each sample. Six outliers were removed as determined by deviation of the summed intensity of all internal standards, or the summed intensity of all annotated known compounds by more than two standard deviations in either positive or negative mode ionization analysis.

Calculation of ANOVA was carried out for all features comparing genotypes (n=7, 8 or 9), and multiple comparisons were accounted for using the Bonferroni correction accounting for 4152 tests conducted. Calculation of p-values for Student T-Test was carried out by comparing the peak heights of a single genotype to the relevant control genotype as noted (Table S3**)**. PLS-DA analysis was calculated using only known annotated metabolites, and not unknown features. Custom R scripts were used to perform these statistical analyses.

### Supplemental Tables

#### Table S1. Panel of 94 phospho-substrates used in HAD activity screens

| **Substrate** | **Abbreviation** | **Substrate** | **Abbreviation** |
|---|---|---|---|
| 2',5'-Adenosine diphosphate | 2',5'-ADP | Adenosine 3',5'-diphosphate | PAP |
| 2'-Adenosine monophosphate | 2'AMP | Adenosine 3'-phosphate 5'-phosphosulfate | PAPS |
| 2'-Cytidine monophosphate | 2'CMP | Phospho(enol)pyruvate | PEP |
| 2-Phospho-ascorbate | 2P-Ascorbate | Pyridoxal 5'-phosphate | PLP |
| 2-Deoxyglucose 6-phosphate | dGlc-6P | Phosphono-acetate | Po-acetate |
| 2-Deoxyribose 5-phosphate | dRib-5P | Phosphono-formate | Po-formate |
| 3'-Adenosine monophosphate | 3'AMP | *N*-(Phosphonomethyl)glycine | Po-methyl-Gly |
| 3'-Cytidine monophosphate | 3'CMP | Phosphorylethanolamine | P-ethanolamine |
| 3-Phosphoglyceraldehyde | 3-PGA | Phytic acid | Phytate |
| 3-Phosphoglycerate | 3P-Glycerate | Polyphosphate | Poly-P |

| | | | |
|---|---|---|---|
| 6-Phosphogluconate | 6P-Gluc | Inorganic pyrophosphate | Ppi |
| Adenosine diphosphate | ADP | Ribose 5-phosphate | Rib-5P |
| Adenosine monophosphate | AMP | Ribulose-1,5-diphosphate | Ru-1,5-P2 |
| Adenosine monophosphoramidate | AMP-ram | Sucrose 6-phosphate | Sucrose-6P |
| Adenosine triphosphate | ATP | Thymidine diphosphate | TDP |
| *N*,*N*-Bis(phosphonomethyl)glycine | BPMG | Thymidine triphosphate | TTP |
| Cytidine diphosphate | CDP | Thiamine monophosphate | Thiamine-P |
| Cytidine monophosphate | CMP | Thiamine pyrophosphate | Thiamine-PP |
| Cytidine triphosphate | CTP | Trehalose 6-phosphate | Trehalose-6P |
| Coenzyme A | CoA | Uridine diphosphate | UDP |
| Dihydroxyacetone phosphate | DHAP | Uridine monophosphate | UMP |
| Erythrose 4-phosphate | Eryth-4P | Uridine monophosphate-morpholidate | UMP-mor |
| Riboflavin 5'-monophosphate | FMN | Uridine triphosphate | UTP |
| Fructose-1,6-diphosphate | Fru-1,6 diP | Xanthosine monophosphate | XMP |
| Fructose 1-phosphate | Fru-1P | α-D-Glucosamine 1-phosphate | a-D-Glcn 1P |
| Fructose 6-phosphate | Fru-6P | α-Glucose 1-phosphate | a-Glc-1P |
| Guanosine diphosphate | GDP | β-Glucose 1-phosphate | b-Glc-1P |
| Guanosine monophosphate | GMP | Glucose 6-phosphate | Glc-6P |
| GMP-morpholidate | GMP-mor | Deoxyadenosine diphosphate | dADP |
| Guanosine triphosphate | GTP | Deoxyadenosine monophosphate | dAMP |
| Galactose 1-phosphate | Gal-1P | Deoxyadenosine triphosphate | dATP |
| Glucosamine 6-phosphate | Glcn-6P | Deoxycytidine diphosphate | dCDP |
| Glucose-1,6-diphosphate | Glucose-1,6P2 | Deoxycytidine monophosphate | dCMP |
| Glycerol 1-phosphate | Glyc-1P | Deoxycytidine triphosphate | dCTP |
| Glycerol 2-phosphate | Glyc-2P | Deoxyguanidine diphosphate | dGDP |
| Glycerol 3-phosphate | Glyc-3P | Deoxyguanidine monophosphate | dGMP |
| Glyphosate | Glyphosate | Deoxyguanidine triphosphate | dGTP |
| Inosine diphosphate | IDP | Deoxyinosine monophosphate | DIMP |
| Inosine monophosphate | IMP | Deoxyinosine triphosphate | DITP |
| Inosine triphosphate | ITP | Deoxythymidine monophosphate | dTMP |
| L-2-Phosphoglycerate | 2P-Gly | Deoxyuridine monophosphate | dUMP |
| Lactose 1-phosphate | Lactose-1P | Deoxyuridine triphosphate | dUTP |
| Mannose 1-phosphate | Man-1P | Phosphorylcholine | P-Cho |
| Mannose 6-phosphate | Man-6P | Phosphorylserine | P-Ser |
| *N*-Acetyl-α-D-glucosamine-1-phosphate | NAGlcn-1P | Phospho-threonine | P-Thr |
| *N*-Acetyl-α-D-glucosamine-6-phosphate | NAGlcn-6P | Phospho-tyrosine | P-Tyr |
| β-Nicotinamide adenine dinucleotide phosphate | NADP | *p*-Nitrophenylphosphate | pNPP |

**Table S2. LC-MS/MS metabolomic data processing parameters for open source software**

| MSDIAL v. 3.70 settings | HILIC Positive mode | HILIC Negative mode |
|---|---|---|
| Data collection MS1 tolerance | 0.01Da | 0.01Da |
| Data collection MS2 tolerance | 0.025 Da | 0.025 Da |
| Mass Range | 60-900 Da | 60-900 Da |
| smoothing level | 3 | 3 |
| Minimum peak width | 10 | 10 |
| Minimum peak height | 100000 | 100000 |
| mass slice width | 0.1 | 0.1 |
| MS/MS matching: MS1 accurate mass tolerance | 0.005 Da | 0.005 Da |
| MS/MS matching: MS2 accurate mass tolerance | 0.05 Da | 0.05 Da |
| m/z & RT match time tolerance | 0.15 minutes | 0.1 minutes |
| m/z & RT match mass tolerance | 0.005 Da | 0.005 Da |
| MS/MS libraries used for analysis | MoNA, NIST17 | MoNA, NIST17 |

**MS-DIAL version 3.70.**

**Table S3.** Statistical comparison of genotypes (n=7, 8 or 9) using a 1-way Analysis of Variance with p-value corrected for multiple comparisons using the Bonferroni correction. T-tests were run by comparing one genotype as listed, to either 3A for mutants, or 230A against 230H. No signals were detected in 3A for 2-deoxyinosine-5-monophosphate which making this test infeasible

| Metabolite Name | 1-way ANOVA (p-value) | 1-way ANOVA (p-value corrected for multiple comparisons) | T-test (groups compared) | T-test (p-value) |
|---|---|---|---|---|
| Uridine diphosphate hexose | 2.72E-32 | 1.13E-28 | not tested | |
| beta-Glycerophosphate | 5.32E-26 | 2.21E-22 | 728 vs. 3A | 7.06E-06 |
| Thiamine cation | 3.71E-14 | 1.54E-10 | 66 vs. 3A | 1.64E-08 |
| PG 30:0 | 4.03E-18 | 1.67E-14 | 230A vs. 230H | 4.04E-06 |
| Deoxyuridine monophosphate | 4.28E-62 | 1.78E-58 | 66 vs. 3A | 7.40E-15 |
| PG 31:0 | 2.24E-18 | 9.28E-15 | 230A vs. 230H | 4.76E-08 |
| Inosine-5-monophosphate | 2.83E-40 | 1.17E-36 | 66 vs. 3A | 1.00E-07 |
| Oleoyl-alpha-lysophosphatidic acid | 6.64E-43 | 2.76E-39 | 728 vs. 3A | 4.96E-09 |
| Hexadecyl-octadecenoyl-glycero-phosphocholine | 1.00E-04 | 5.50E-01 | not tested | |
| 1-Palmitoylglycerol | 1.76E-37 | 7.31E-34 | 728 vs. 3A | 8.64E-07 |
| 2-Deoxyinosine-5-monophosphate | 4.10E-26 | 1.70E-22 | 66 vs. 3A not possible | not detected in 3A |
| Cytidine | 4.98E-16 | 2.07E-12 | 230A vs. 230H | 5.36E-07 |
| PA 34:1 | 2.60E-13 | 1.08E-09 | not tested | |
| 9-Oxo-10,12-octadecadienoic acid | 6.85E-19 | 2.84E-15 | not tested | |
| Fructose-1-phosphate | 5.14E-24 | 2.14E-20 | not tested | |

**Table S4.** Strains and plasmids used in this study.

| Strain | Phenotype, genotype and/or description[a] | Ref/Source |
|---|---|---|
| *E. coli* | | |
| GC10 | F- mcrA Δ(mrr-hsdRMS-mcrBC) Δ80dlacZΔM15 ΔlacX74 endA1 recA1 Δ(ara, leu)7697 araD139 galU galK nupG rpsL λ- T1R | GeneChoice, Inc |
| BW25113 | F-, Δ(araD-araB)567ΔlacZ4787(::rrnB-3) λ-, rph-1, Δ(rhaD-rhaB)568hsdR514 | (12) |
| **MG1655** | F- lambda- ilvG- rfb-50 rph-1 | *E. coli* Genetic Stock Center |
| **BL21(DE3)** | *ompT gal dcm lon hsdS*$_B$(*r*$_B^-$*m*$_B^-$) λ(DE3 [*lacI lacUV5-T7p07 ind1 sam7 nin5*]) [*malB*$^+$]$_{K\text{-}12}$(λ$^S$) | Novagen |
| **BL21(DE3) pLysS** | *ompT gal dcm lon hsdS*$_B$(*r*$_B^-$*m*$_B^-$) λ(DE3 [*lacI lacUV5-T7p07 ind1 sam7 nin5*]) [*malB*$^+$]$_{K\text{-}12}$(λ$^S$) pLysS[*T7p20 ori*$_{p15A}$](Cm$^R$) | (13) |
| **BL21 CodonPlus (DE3)-RIPL** | *E. coli* B F– *ompT hsdS*$_B$(*r*$_B^-$*m*$_B^-$) dcm+ Tet$^R$ gal λ (DE3) *endA* Hte [argU ileY leuW Camr] | Stratagene |
| **JW1950** | BW25113 ΔhchA::Kan$^R$ | (12) |
| **JW5057** | BW25113 ΔyajL::Kan$^R$ | (12) |
| **JW0097-KC** | BW25113 ΔmutT::Kan$^R$ | (12) |
| **JW0097-AM** | W3110 ASKA-mutT | (14) |
| **DCH1198** | BW25113 ΔyajL::Sin FRT | This study |
| **DCH1204** | BW25113 ΔyajL::Sin FRT ΔhchA::Kan$^R$ | This study |
| **DCH1152** | MG1655 ΔygfA::Sin FRT | This study |
| **DCH1202** | JW2752 pBAD24-380 | This study |

| | | |
|---|---|---|
| **DCH1215** | JW0097 pBAD24 | This study |
| **DCH1180** | JW0097 ASKA-mutT | This study |
| **DCH1201** | JW0097 pBAD24-380 | This study |
| **DCH1277** | JW0097 pBAD24-nadD$_M$ | This study |
| **DCH1278** | JW0097 pBAD24-yqeK$_M$ | This study |
| **DCH1166** | JW5057 pUC19 | This study |
| **DCH1171** | JW5057 pUC19-400 | This study |
| **DCH1220** | DCH1204 pUC19 | This study |
| **DCH1233** | DCH1204 pUC19-400 | This study |
| ***Mycoplasma*** | | |
| **JCVI-syn1.0** | | (15) |
| **JCVI-syn3A** | | (5) |
| **Syn3A_Δ66+Puro** | JCVI-syn3A *ΔJCVISYN3A_0066*::Pur$^R$ | This study |
| **Syn3A_Δ77+Puro** | JCVI-syn3A *ΔJCVISYN3A_0077*::Pur$^R$ | This study |
| **Syn3A_Δ728+Puro** | JCVI-syn3A *ΔJCVISYN3A_0728*::Pur$^R$ | This study |
| **Syn3A_Δ907+Puro** | JCVI-syn3A *ΔJCVISYN3A_0907*::Pur$^R$ | This study |
| **Syn3A_0380 H230A** | JCVI-syn3A *JCVISYN3A_0380*: H230A | This study |
| **Syn3A_0380 H230H(C-T)** | JCVI-syn3A *JCVISYN3A_0380*: H230H (C-T) | This study |
| **Plasmids** | | |
| **pUC19** | Amp$^R$, ColE1, IPTG inducible promoter | (16) |
| **pBAD24** | Amp$^R$, ColE1, arabinose inducible promoter | (14) |
| **pUC57** | InfoAmp$^R$, pMB1, IPTG inducible promoter | |

| | | |
|---|---|---|
| **pET28a** | pBR332 lacIpT7 Kan$^R$ His6tag expression vector | Novagen, Inc |
| **pCP20** | Yeast FLP recombinase Cm$^R$ Amp$^R$ oriR101 w/repA101ts | (17) |
| **ASKA-mutT** | pCA24N mutT Cm$^R$ | (14) |
| **pUC19-400** | pUC19 optimized JCVISYN3A_0400 | This study |
| **pUC19-443** | pUC19 optimized JCVISYN3A_0443 | This study |
| **pUC19-887** | pUC19 optimized JCVISYN3A_0887 | This study |
| **pUC57-380** | pUC57 optimized JCVISYN3A_0380 | This study |
| **pET28-887** | pET28 optimized JCVISYN3A_0887 HisTag C-ter | This study |
| **pBAD24-380** | pBAD24 optimized JCVISYN3A_0380 | This study |
| **pET28-066** | pET28 optimized JCVISYN3A_0066 HisTag C-ter | This study |
| **pET28-077** | pET28 optimized JCVISYN3A_0077 HisTag C-ter | This study |
| **pET28-728** | pET28 optimized JCVISYN3A_0728 HisTag C-ter | This study |
| **pET28-907** | pET28 optimized JCVISYN3A_0907 HisTag C-ter | This study |
| **pET28-380** | pET28 optimized JCVISYN3A_0380 HisTag N-ter | This study |
| **pET28-380H230A** | pET28 optimized JCVISYN3A_0380 HisTag N-ter with mutated His230 | This study |
| **pET28a-nadD$_{Bs}$** | pET28 NadD$_{Bs}$optimized JCVISYN3A_0907NadD$_{Bs}$ HisTag C-ter | This study |
| **pET28a-yqeK$_M$** | pET28 optimized JCVISYN3A_0380 ykeK domain only | This study |
| **pBAD24-nadD$_M$** | pBAD24 optimized JCVISYN3A_0380 nadD domain only | This study |
| **pBAD24-yqeK$_M$** | pBAD24 optimized JCVISYN3A_0380 yqeK domain only | This study |
| **pET15a-400** | pET15a optimized JCVISYN3A_0400 HisTag | This study |

| | | |
|---|---|---|
| **Pmod2*lox*puro*lox*-cre-sp** | Plasmid for insertion of genes into loxP landing pads in JCVI-syn3B | GenBank: MN982903.1 and (18) |

[a]Abbreviations : Kan$^R$, kanamycinkanamycinekanamycin resistance; Cm$^R$, chloramphenicol resistance; Amp$^R$, ampicillin resistance; Pur$^R$ ,puromycin resistance; C-ter, C-terminal; N-ter, N-terminal.
